# The effects of environmental complexity on microbial community yield and structure

**DOI:** 10.1101/2020.07.09.195404

**Authors:** Alan R. Pacheco, Daniel Segrè

## Abstract

Environmental composition is a major, though poorly understood determinant of microbiome dynamics. Here we ask whether general principles govern how key microbial community properties, i.e. yield and diversity, scale with the number of environmental molecules. By assembling hundreds of synthetic consortia, we found that community yield remains constant as a function of environmental complexity, in agreement with additive expectations of an idealized model. However, taxonomic diversity is much lower than expected. By quantifying this deviation with a metric for epistatic interactions between environments, we uncovered simple ecological rules that govern how communities respond nonlinearly to the coupling of different nutrient sets. Our results demonstrate that environmental complexity alone is not sufficient for maintaining microbiome diversity, and provide practical guidance for designing and controlling microbial communities.

## Introduction

Changes in environmental molecules can dramatically alter microbial community properties, as observed in dietary shifts in humans^1,2^ or nutrient fluctuations in marine or terrestrial ecosystems^3,4^. Moreover, in the nascent field of synthetic ecology, a central unresolved challenge is how to design communities with specific functions by engineering their molecular environments^5–7^. Despite its importance, we lack a generalized understanding of how environmental complexity, i.e. the number of different substrates, affects microbial ecosystems. Although classical ecological theories based on competitive exclusion and niche partitioning would suggest greater biodiversity in more complex environments^8,9^, factors such as organism-specific metabolic capabilities^10,11^ and interspecies interactions^12,13^ can cause significant deviations from this expectation. More recent studies exploring environmental complexity have reported conflicting results on its role in modulating growth yields and biodiversity^14–17^, motivating the need for a systematic exploration of its effects in well-controlled conditions. Here, we quantify how the yield and diversity of over 280 synthetic communities scale with increasing environmental complexity, examining how these properties differ from expectations based on those in simpler environments. Through a quantitative analysis by which we compare our data with mathematical models, we reveal that these scaling relationships can be explained by a set of ecological rules for combining environments, with implications for the ecology of natural and engineered microbiomes.

## Results

### Assembly of multispecies communities and combinatorial nutrient conditions

We first designed microcosms with varying degrees of initial taxonomic and substrate complexity (Figure 1a-d). Based on an experiment that assessed the nutrient utilization capabilities of various bacterial species (Supplementary Figure 1, Supplementary Figure 2, Supplementary Table 1, Supplementary Table 2), we selected 13 organisms and 32 carbon sources intended to maximize taxonomic variability across environments. These organisms, which are not representative of any particular biome, were also chosen as they can be readily cultured individually and introduced into combinatorial environments in a controlled way. We generated increasingly complex combinations of our 32 nutrients in a hierarchical manner, so that we could quantitatively compare the effects of higher-order combinations with those of simpler ones (Figure 1d). Additionally, each environment contained the same amount of carbon irrespective of nutrient complexity (Figure 1b), enabling us to specifically assess the impact of increased resource heterogeneity. Our organisms were inoculated into these environments at equal amounts (Supplementary Table 3, Supplementary Table 4), and the resultant cultures were grown and passaged into fresh media at rates informed by pilot experiments in order to maximize the chance of each having consumed the provided substrates and reached a stable composition (Supplementary Figure 3, Supplementary Figure 4b). In total, variants of this procedure were applied to generate 282 unique community-environment pairings (Supplementary Table 5).

**Figure 1.**
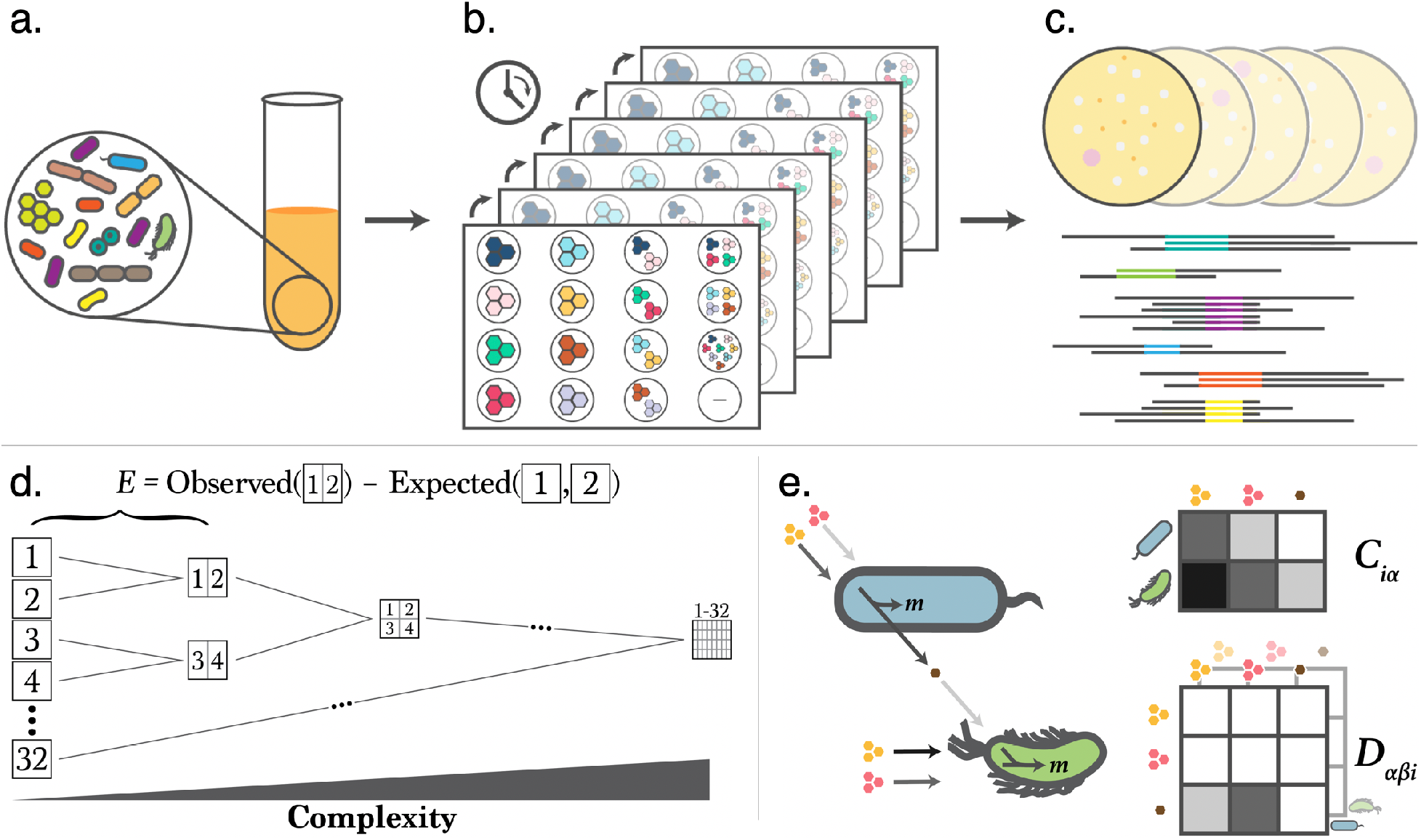
Experimental schematic for testing microbial community responses to environmental complexity. **(a).** Communities were assembled by combining a defined number of organisms (Supplementary Figure 1a, Supplementary Table 1) at equal ratios. **(b).** These mixed cultures were then inoculated into deep-well plates containing a minimal medium plus equimolar combinations of up to 32 carbon sources (Supplementary Figure 1b, Supplementary Table 2, Supplementary Table 3, Supplementary Table 4). The communities were either grown in batch or diluted into fresh media over the course of several days (Supplementary Table 5). **(c).** Biomass yields were then assessed using a spectrophotometer and composition was determined using either agar plating or 16s sequencing. **(d).** Experiments are designed such that community phenotypes in more complex environments can be directly compared to simpler environments containing the same nutrients. Simpler environments are used to generate expectations of phenotype in more complex compositions. **(e).** Monod consumer resource modeling framework. Resources (pink and yellow) are utilized by organisms according to a stoichiometric matrix *C* and converted into biomass *m*. In this example, the bottom green organism is able to utilize both resources more efficiently than the top blue one as denoted in the shades in the *C* matrix and of the nutrient-organism arrows. The blue organism converts both the pink and yellow nutrients into a brown metabolite according to the species-specific stoichiometric matrix *D*. This secreted metabolite cannot be consumed by the blue organism in this example, but it can be utilized by the green organism as indicated by the *C* matrix and the arrows.

### Measuring synthetic community yield as a function of environmental complexity

We initially asked whether and how the yield of a community as a whole varies with increasing environmental complexity. To generate an expectation of this effect, we applied a consumer resource model (CRM, Figure 1e)^18^ to a statistical ensemble of simulated communities. This model predicted that, on average, community yield does not change significantly with environmental complexity (Supplementary Figure 5). Indeed, despite comprising a diverse set of organisms on heterogeneous nutrient combinations, our *in vitro* 13-species community data closely matched this expectation (Figure 2a). To further explore possible deviations from this overall effect, we established a metric that quantifies how much the yield *Y* on a specific composite environment differs from the expectation based on its constituent environments (Figure 1d). This ‘epistasis’ metric *E*_*Y*_, similar to those used to describe nonlinearities between genetic^19^ or environmental^20^ perturbations, is defined as the difference between the observed and expected yield on the combination of two environments *A* and *B*, i.e., *E*_*Y*_ = *Y*(*AB*) − (*Y*(*A*) + *Y*(*B*))/2. An *E*_*Y*_ value of zero would thus reflect the naïve assumption that, since all environments contain the same total amount of carbon, the yield in a complex environment should be the same as the combination of yields on its corresponding simpler environments.

**Figure 2.**
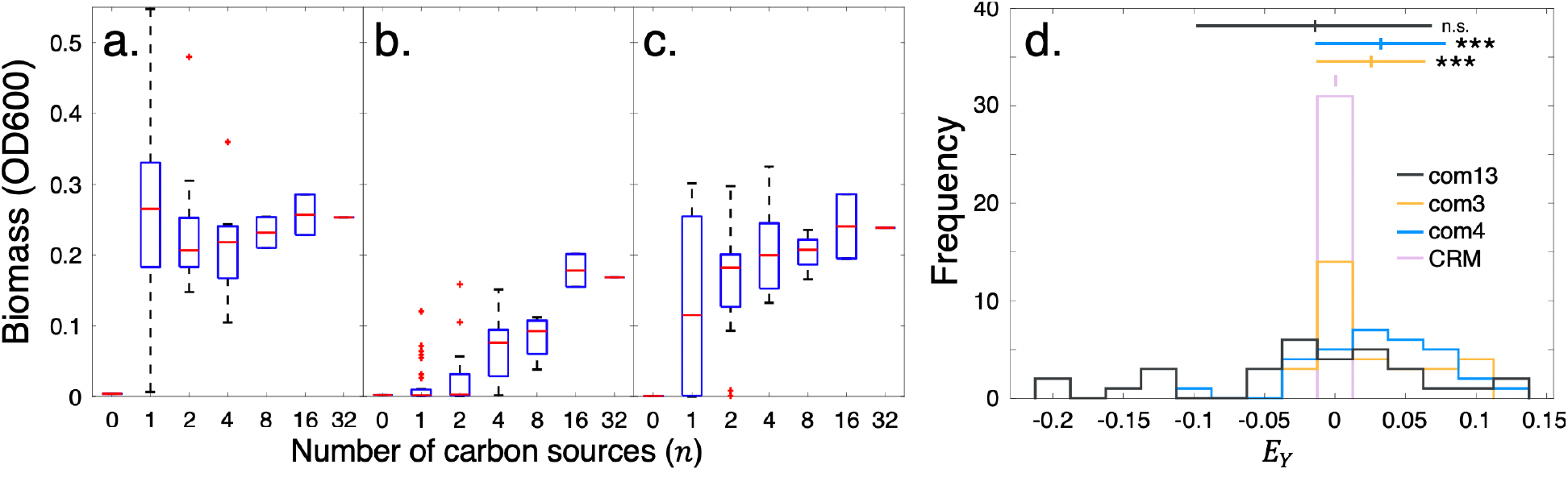
Changes in community yield in response to environmental complexity. **(a-c).** Biomass yields for com13 **(a)**, com3 **(b)**, and com4 **(c)** measured at the experimental endpoint (6 passages at 48-hour frequencies, Supplementary Table 5). Here, the central mark indicates the median, the top and bottom box edges indicate the 25^th^ and 75^th^ percentiles, respectively, the whiskers extend to the most extreme points not considered outliers, and the red ‘+’ symbols indicate outliers plotted individually. Sample sizes are outlined in Supplementary Table 3. **(d).** Distribution of yield epistasis *E*_*Y*_ for all three communities and simulated communities predicted using a consumer resource model (CRM). The simulated communities are composed of 13 organisms whose nutrient utilization capabilities were randomly sampled 50 times from a uniform distribution. Upper bars denote mean and standard deviation. Significance levels are calculated between each community distribution and that of the CRM using a paired t-test, and are indicated by (***) *p* < 0.001 (*p* = 0.84 for com13, 3.6 × 10^−4^ for com3, and 2.1 × 10^−4^ for com4).

The distributions of yield epistasis scores *E*_*Y*_ for our 13-species community (referred to as com13) and for the CRM simulations were centered at zero (Figure 2d), confirming that our experiment and the corresponding model match our expectation of yield additivity. Despite this overall behavior, various nutrient combinations in our experiments resulted in significant deviations from *E*_*Y*_ = 0 (Supplementary Figure 6). For example, the community cultured on the combination of D-glcNAc and D-galacturonate displayed an *E*_*Y*_ value of 0.13 (2*σ*), indicating improved growth on this more complex composition than on the individual nutrients. Conversely, the combination of D-glucose and D-sorbitol resulted in an *E*_*Y*_ score of −0.19 (3*σ*), suggesting that the community might be displaying reduced efficiencies in using one substrate in the presence of another, representing a type of ‘resource interference’ previously observed experimentally^14,21^.

While our 13-species community matched the expectation that yields are additive on average, we realized that this property could reflect at least two distinct underlying effects. On the one hand, additivity might be a property of the metabolism of individual organisms, which is propagated to the community level when organisms are cultured together. On the other hand, it could be an ecological property that emerges only in complex communities. To address this question, we compared the 13-species community results with those of similar experiments performed on smaller (3- and 4-species) consortia, as well as on individual organisms. Surprisingly, the yields observed in these experiments deviated from the constant overall yield observed for the complex communities, instead increasing with environmental complexity (Figure 2b, c, Supplementary Figure 7, Supplementary Figure 8). Correspondingly, for both of the small communities and some of the individual organisms, the distribution of *E*_*Y*_ was significantly skewed in the positive direction (Figure 2d, Supplementary Figure 8), suggesting that the additivity of yields on combined environments may not hold for individual organisms and small communities. This effect may be due to metabolic nonlinearities associated with each individual organism, which dominate the scaling of yield for small communities. However, these nonlinearities may dampen upon reaching a certain threshold of community complexity, giving rise to the flat yield scaling observed for com13 (Figure 2a, d). This would imply that the average yield additivity emerges as an ecological property, which may balance the metabolic synergy present in individual organisms (Supplementary Text).

### Environmental complexity is not necessary for maintenance of taxonomic diversity

Our analysis has so far focused on a single collective trait of microbial communities, i.e. the total yield, but has not provided insight into how environmental complexity affects the balance between different organisms and the ensuing community structure. We thus used 16s amplicon sequencing to measure the endpoint taxonomic distributions of two 13-species communities under increasingly complex environments (com13, com13a, Supplementary Table 5). This analysis revealed considerable variation across different environments (Figure 3a, Supplementary Figure 9) and high degrees of consistency across replicates and experiments irrespective of environmental complexity (Supplementary Figure 10, Supplementary Figure 11), suggesting that the assembly patterns of these communities are largely deterministic based on nutrient composition. To more deeply analyze the contributions of defined nutrient sets to specific community structures, we applied a clustering analysis that yielded an environment-phenotype mapping spanning our entire dataset (Supplementary Figure 21). This mapping demonstrated how distinct – and often unrelated – environments can nonintuitively result in similar taxonomic compositions, which reflect previously-identified family-level functional relationships in natural microbiomes (Supplementary Text, Supplementary Figure 22)^12^.

**Figure 3.**
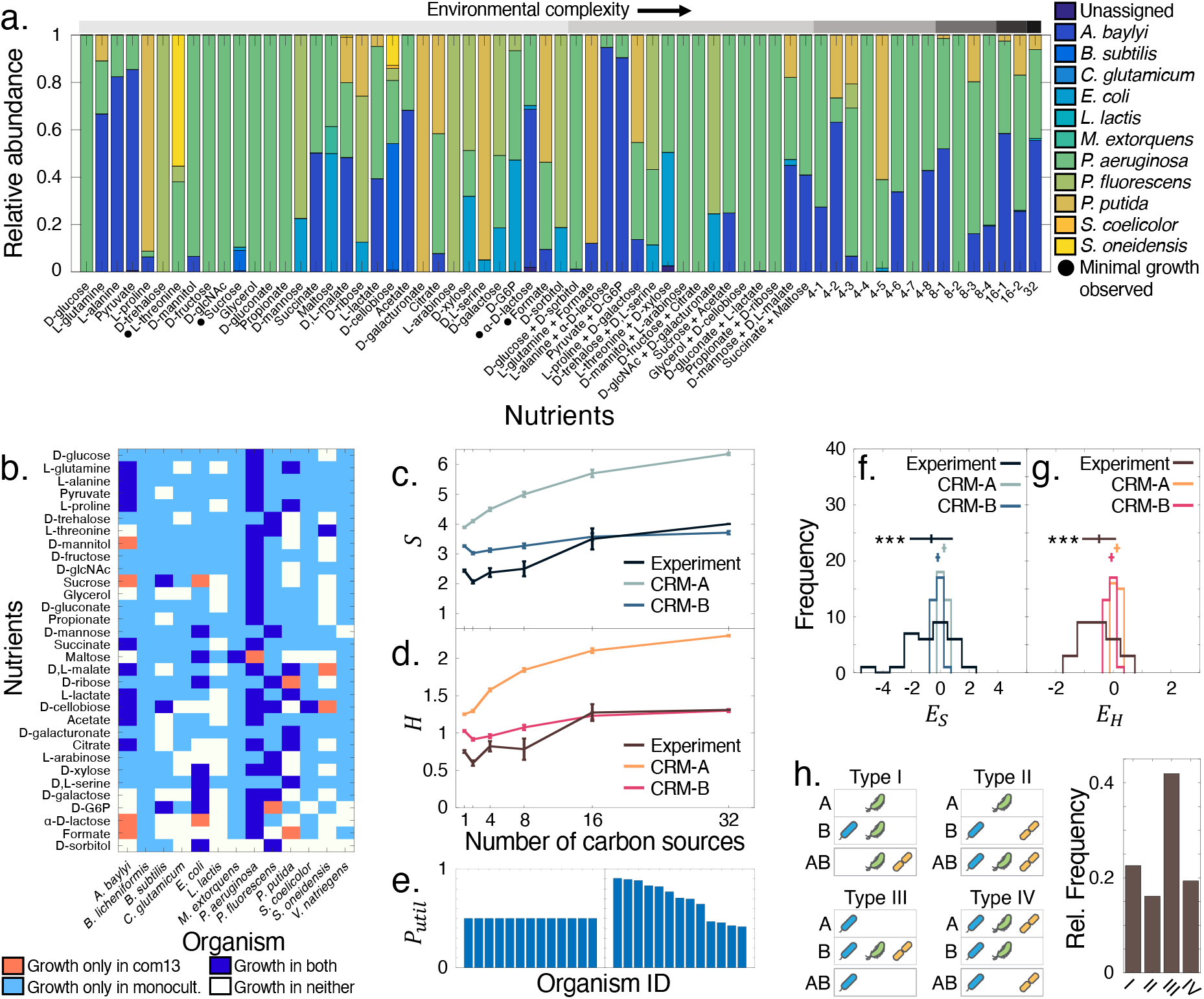
Endpoint taxonomic properties of 13-species community in up to 32 carbon sources (com13). **(a).** Mean species relative abundances. Environments with more than two nutrients are abbreviated (e.g. condition 4-1 contains the carbon sources in the first two 2-nutrient conditions, etc.). For complete environmental compositions see Supplementary Table 3. Gray circles indicate community growth below OD600 0.05. Initial compositions and compositions across all replicates are found in Supplementary Figure 11a. **(b).** Species-specific differences in growth between monoculture (Biolog assay, Supplementary Figure 2a) and single-carbon source community contexts. **(c, d).** Comparison of observed community species richness *S* **(c)** and Shannon entropy *H* **(d)** with phenotypes predicted by consumer resource models. The nutrient utilization capabilities of simulated organisms are either the same on average (CRM-A) or variable, allowing for generalists and specialists (CRM-B). Error bars indicate s.e.m. No significant increases in *S* or *H* were identified when comparing the single-nutrient cases to the 32-nutrient cases (one-tailed paired t-test *p* = 0.107 for *S* and 0.180 for *H*). **(e).** Representation of the fraction of nutrients usable by each organism (*P*_*util*_). Organisms are sorted by decreasing *P*_*util*_. Left: each organism has the same fixed probability of consuming a given nutrient. Right: this probability varies for each organism, determined by the fraction of nutrients that were consumed by each organism in our initial phenotypic assay (Supplementary Figure 2a). **(f, g).** Distributions of species richness epistasis (*E*_*S*_, **f**) and Shannon entropy epistasis (*E*_*H*_, **g**) scores for experiment and CRM. Upper bars denote mean and standard deviation. Significance values are calculated against the distributions for CRM-A and are indicated by (***) *p* < 0.001 (*p* = 2.0 × 10^−4^ for *E*_*S*_ and 1.6× 10^−7^ for *E*_*H*_). **(h).** Schematic and prevalence of different epistasis types. Illustrations are representative examples. Type I: The environment AB results in the presence of an organism not observed in either environment A or B; Type II: AB results in the union of organisms from A and B; Type III: AB contains only the organisms from the lowest-diversity environment; Type IV: AB results in a more complex loss of diversity.

Though the overall species abundance distributions in our communities were similar to those of natural microbiota^22–24^ (Supplementary Figure 12), we noticed that very few organisms out of the original 13 persisted in any single environment (Figure 3a, Supplementary Figure 9). To determine the possible cause of these losses in diversity, we examined the taxonomic compositions of com13 in single carbon sources, and found that many organisms did not persist in a community context despite being able to utilize a given substrate in monoculture (Figure 3b). This discrepancy was particularly striking in the D-glucose condition, in which, despite all but one organism being able to metabolize this carbon source, only *P. aeruginosa* remained. In addition, organisms that displayed a wide range of metabolic capabilities in monoculture, such as *C. glutamicum*, were not observed at all in the single-nutrient environments, suggesting that the structure of our communities was largely driven by competition. In support of this reasoning, we also found generally poor correlations between the yields reached by the organisms in monoculture and in com13 (Supplementary Figure 13), indicating that monoculture resource utilization patterns are not necessarily predictive of how an organism will behave in a community. Despite the prevalence of interspecies competition, there were instances in which organisms unable to grow in monoculture on a given nutrient did survive in the same nutrients in a community context (Figure 3b). For example, the community grown on D-ribose contained *P. putida* despite its inability to utilize this substrate (Supplementary Figure 2b). A possible explanation is that *P. fluorescens*, which was present in high abundance, secreted organic acids that sustained *P. putida* after catabolizing D-ribose into glyceraldehyde 3-phosphate. Indeed, such metabolic transformations are well-documented in strains of *P. fluorescens* that share the relevant genes with our strain^25–27^.

The availability of hierarchical nutrient combinations gave us the opportunity to extend our analysis of diversity to multi-nutrient conditions and ask whether, beyond anecdotal cases, general principles seem to govern the scaling of diversity at increasing environmental complexity. We thus first used our 16s data to calculate the species richness *S* and Shannon entropy *H* values of our communities at each degree of environmental complexity – from 1 to 32 carbon sources. Our initial expectation that more carbon sources would create more niches and therefore lead to higher diversity was clearly contradicted by the data, as neither diversity metric increased significantly as a function of environmental complexity (Figure 3c, d). In fact, some single-carbon source conditions resulted in greater diversity than other more complex environments, suggesting that the number of substrates is not a key determining factor of taxonomic diversity for these communities. Moreover, species co-occurrence patterns that we observed in single-nutrient environments were not preserved in more complex settings (Supplementary Figure 14).

In order to more systematically assess the effects of combinations of environments on diversity, we defined epistasis metrics *E*_*S*_ and *E*_*H*_ for species richness and Shannon entropy, respectively. Like our epistasis scores for yield, these metrics quantify changes in taxonomic diversity based on expectations of these quantities on simpler environments:

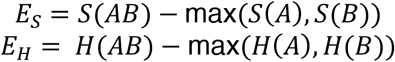

These scores draw from expectations of a lower bound for *S* and *H* based on the intuition that a community on a more complex environment *AB* should be at least as taxonomically diverse as that on *A* or *B*. By computing these epistasis values for community compositions simulated using our consumer resource model, we found that the predicted distributions of *E*_*S*_ and *E*_*H*_ were both centered at zero (Figure 3f, g), confirming that our definition of epistasis constitutes a reasonable baseline to which we could compare our experimental results. In contrast to this basic expectation, but consistent with the low levels of diversity observed experimentally, the distributions of both scores for our *in vitro* 13-species communities were significantly skewed to the left (*E*_*S*_ = −0.65 ± 1.47, *E*_*H*_ = −0.50 ± 0.58; Figure 3f, g). In other words, our experiments revealed the pervasive presence of negative epistasis in how diversity behaves upon mixing two sets of resources.

To better understand the causes underlying this phenomenon, we examined how the taxonomic compositions exhibited in individual environments translated to those in combinations of environments. We found that the taxonomic outcomes of combining two nutrient sets could be classified into four basic types (Figure 3h, Supplementary Data 1). Interestingly, in about 20% of the cases displayed, the combined environment resulted in the appearance of one or more organisms that had not grown on the individual environments (Type I), suggesting the presence of beneficial interspecies interactions. However, the most common pattern emerging from our data was the dominance of the least diverse constituent environment (Type III, accounting for 40% of the cases). All instances of this dominance, which resembles complete buffering epistasis^19^, were associated with strongly negative values of *E*_*S*_, accounting in large part for the overall negative bias of the distribution.

The prevalence of the Type III pattern highlighted that even complex combinations of nutrients often lead to the dominance of a single organism (Supplementary Data 1). We thus sought to determine whether this observation could be explained by explicitly considering the nutrient use capabilities of our organisms. To do this, we applied our consumer resource model to simulate two sets of 13-species communities: one based on the naive assumption that all organisms consume the same number of usable nutrients on average (CRM-A), and another in which the proportion of nutrients usable by each organism was based on our initial experimental phenotypic assay (Supplementary Figure 2a), thereby reflecting the presence of metabolic generalists and specialists (CRM-B, Figure 3e). We found that, while species richness and Shannon entropy were predicted to increase with environmental complexity in the CRM-A communities (*S* reaching a maximum of ~7 coexisting species), they remained relatively flat in CRM-B communities (*S* reaching a maximum of ~3 coexisting species, a value very similar to experimental observations (Figure 3c)). Predicted epistasis scores were also negatively skewed in CRM-B (Figure 3f, g), suggesting that reduced taxonomic diversity in complex environments could be the outcome of competition in communities with uneven metabolic capabilities (Supplementary Figure 15). The important role of generalists in determining the final composition of a community was also confirmed by more detailed analyses of smaller communities, which illustrated that metabolic exchange could slightly dampen losses in taxonomic diversity (Supplementary Text).

## Discussion

Disentangling how multispecies microbial communities grow on mixtures of substrates remains highly challenging. Though our simplified experimental system is still far from the complexity of natural microbiomes, it captures properties that go beyond those observable in small artificial consortia. In particular, we identified a simple additive principle that explains how average total yields remain invariant with increasing environmental complexity – a consequence of all available resources being efficiently utilized given enough organisms with varied metabolic capabilities. Though one could expect this behavior to arise in communities well adapted to a specific environment, it is surprising that it also emerged in our synthetic consortia composed of organisms from different biomes grown on artificial combinations of carbon sources. The notion that synthetic communities may spontaneously converge to states resembling natural communities is also corroborated by the similarity of our taxonomic abundance distributions to those observed for a number of complex microbiota (Supplementary Text, Supplementary Figure 12). A closer analysis of the compositions of our communities also revealed that increased environmental complexity does not guarantee greater taxonomic diversity beyond that already possible on individual nutrients^12^. This result underscores the dependence of biodiversity on an interplay of features, such an appropriate balance of generalists and specialists and the existence of evolved interdependencies^17,28,29^. An improved understanding of the rules that govern how nutrient sets combine will therefore be necessary for designing complex communities with desired taxonomic properties, as well as for the generation of phenomenological laws and multiscale models that can shed light on the role of communities in host-associated microbiomes and biogeochemical cycles^30–34^.

## Methods

### Selection and initial metabolic profiling of organisms

In order to maximize the chance of obtaining communities with diverse taxonomic profiles from different environmental compositions, the organisms selected were drawn from a number of bacterial taxa known to employ varying metabolic strategies. In addition, given the growing relevance of synthetic microbial communities to industrial and biotechnological applications^35–38^, we chose to employ bacterial species that have previously been used as model organisms and have well-characterized metabolic capabilities. This criterion, paired with the availability of flux-balance models associated with a majority of these organisms, allows us to explore the metabolic mechanisms observed in our various experimental conditions with higher confidence. These selection principles resulted in a set of 15 candidate bacterial organisms spanning three bacterial phyla (Actinobacteria, Firmicutes, and Proteobacteria, Supplementary Table 1, Supplementary Figure 1a).

A microtiter plate-based phenotypic assay was used to assess the metabolic capabilities of each of the 15 candidate organisms. Each organism, stored in glycerol at −80°C, was initially grown in 3mL of Miller’s LB broth (Sigma-Aldrich, St. Louis, MO) for 18 hours with shaking at 300 rpm at each organism’s recommended culturing temperature (Supplementary Table 1). To maximize oxygenation of the cultures and prevent biofilm formation, culture tubes were angled at 45° during this initial growth phase. Candidate organism *Streptococcus thermophilus* was found to have produced too little biomass in this time period and was grown for an additional 8 hours. Each culture was then separately washed three times by centrifuging at 8000 rpm for 2 minutes, removing the supernatant, suspending the pellet in 1mL of M9 minimal medium with no carbon source, and vortexing or triturating to homogenize. The cultures were then diluted to OD600 0.5 ± 0.1 as read by a microplate reader (BioTek Instruments, Winooski, VT) and distributed into each well of three PM1 Phenotype MicroArray Plates (Biolog Inc., Hayward, CA) per organism at final OD600 of 0.05. The carbon sources in the PM1 plates (Supplementary Table 2, Supplementary Figure 1b) were resuspended in 150 μl of M9 minimal media prepared from autoclaved M9 salts (BD, Franklin Lakes, NJ) and filter-sterilized MgSO_4_ and CaCl_2_ prior to inoculation. The cultures in each PM1 plate were incubated at each organism’s recommended culturing temperature with shaking at 300rpm for 48 hours. After this growing period, the OD600 of each culture was measured by a microplate reader to quantify growth. To account for evaporation in the outer wells of the plates, which could yield in inflated OD readings, three ‘evaporation control’ plates with no carbon source were inoculated with bacteria at a final OD600 of 0.05 and incubated at 30°C for 48 hours. The averaged OD600 readings of these plates were subtracted from the readings of the bacterial growth plates to correct for evaporation. A one-tailed t-test was performed using these corrected OD600 values to determine significance of growth above the value of the negative controls (p < 0.05). These final growth yields for the 15 candidate organisms are reported in Supplementary Figure 2a, and aggregated analyses of the growth profiles of the organisms are reported in Supplementary Figure 2b-d.

After this initial metabolic profiling, *Streptococcus thermophilus* was not included in any of the subsequent experiments as it displayed too low of a growth rate in the initial overnight growth phase and grew very minimally (no more than OD600 0.2) in fewer than 20% of the carbon sources in the PM1 plate. After inclusion in an initial mixed-culture experiment (com14, Supplementary Table 5), *Salmonella enterica* was also removed from future experiments due to its high levels of growth on all but one of the PM1 plate carbon sources. Its exclusion, meant to prevent its complete dominance in the subsequent mixed-culture experiments, resulted in a final set of 13 bacterial organisms.

For experiments involving a subset of the 13 organisms, the organisms were chosen to ensure they could be differentiated via agar plating. In the 3-species community experiment involving *E. coli*, *M. extorquens*, and *S. oneidensis* in combinations of 5 nutrients (com3a), the organisms were selected based on their easily differentiable colony morphologies (Supplementary Figure 16). In the second 3- and 4-species community experiments (*B. subtilis*, *M. extorquens*, *P. aeruginosa*, and *S. oneidensis (*com3 and com4)), selection was informed by differentiable colony morphology and additional metabolic criteria based on generalist-specialist relationships. Absolute growth yield on the Biolog PM1 plates was also considered, with the goal of including both high- and low- yielding organisms. Therefore, *P. aeruginosa (*high-yield generalist), *B. subtilis* (low-yield specialist), *M. extorquens* (low-yield generalist), and *S. oneidensis* (high-yield specialist) were selected.

### Selection of nutrients and assembly of combinatorial media

The nutrients used in all experiments were selected from the 95 carbon sources contained in the Biolog PM1 Phenotype MicroArray Plate. This plate contains a variety of molecule types such as mono- and disaccharides, sugar alcohols, organic acids, amino acids, and more complex polymers (Supplementary Table 2, Supplementary Figure 1b). Using the metabolic profiling experiments for each individual organism as a basis (Supplementary Figure 2a), different criteria were established to choose the carbon sources used in each experiment depending on the desired complexity of the environment. An overarching criterion was that each experiment contain at least one sugar, one organic acid, and one amino acid to increase the possibility of synergistic interactions between nutrients and nutrient use orthogonality between the organisms.

For the communities grown in 5 carbon sources (i.e. com3a and com13a in D-Glucose, pyruvate, D-glcNAc, L-proline, and L-threonine), the following criteria were applied: D-glucose was selected as it resulted in the highest yield of each of the individual organisms, pyruvate was an organic acid with relatively high yields, D-glcNAc was a more complex sugar that resulted in varying individual growth yields, and L-proline and L-threonine were amino acids that resulted in generally high and low individual species yields, respectively. Communities were grown in all combinations of these five carbon sources (5 conditions of 1 carbon source, 10 of 2, 10 of 3, 5 of 4, and 1 of 5) for a total of 31 unique environmental compositions (Supplementary Table 4).

The carbon sources for the 32-nutrient experiments were selected based on the following criteria: nutrients in which generalists individually displayed low levels of growth but favored at least one specialist (3 nutrients), nutrients that resulted in high-variance in growth yields across species (5 nutrients), and nutrients that resulted in low-variance in growth yields across species (7 nutrients). These criteria were meant to increase the probability of observing more taxonomically diverse communities. The remaining 21 nutrients were selected based on the total species-specific yields they conferred (Supplementary Figure 2a), with higher-yielding nutrients being prioritized. Communities were grown in selected combinations of these 32 carbon sources (32 conditions of 1 carbon source, 16 of 2, 8 of 4, 4 of 8, 2 of 16, and 1 of 32) for a total of 63 unique environmental compositions (Supplementary Table 3). The selected combinations were chosen based on the Biolog biomass yields of the 13 organisms under each carbon source, with the lowest-yielding nutrient (D-sorbitol) being paired with the highest (D-glucose) followed by the second-lowest and second-highest, etc.

Combinatorial media conditions were assembled by resuspending each carbon source in distilled water to stock concentrations of 1.25 mol C/L and filter sterilizing using 0.2 μm membrane filter units (Nalgene, Rochester, NY). Carbon source stock solutions were stored at 4°C for no longer than 30 days. A liquid handling system (Eppendorf, Hamburg, Germany) was used to distribute the individual carbon source stocks in the appropriate combinations in 96-well plates. The combinatorial nutrient stocks were then sterilized with filter plates (Pall Corporation, Port Washington, NY) via centrifugation at 4000rpm for 2 minutes. These were then combined with M9 minimal medium (containing M9 salts, MgSO_4_, CaCl_2_, and no carbon) and filter-sterilized water to final working concentrations of 50 mM C in 96 deep-well plates (USA Scientific, Ocala, FL) for a total volume of 300 μl. This working concentration was selected such that all organisms would not grow beyond the linear range of OD600 for biomass measurements.

### Culturing and quantification of microbial communities in combinatorial media

All communities were assembled using a bottom-up approach, with each organism initially grown separately and diluted to the same starting concentrations before being combined in mixed cultures. For all experiments, each organism was inoculated from a glycerol stock stored at −80°C into 3 mL of LB broth and incubated at 30°C with shaking at 300 rpm for 18 hours. The culture tubes were angled at 45° to prevent biofilm formation and encourage oxygenation of the cultures. The overnight cultures were then washed three times by centrifuging at 8000 rpm for 2 minutes, resuspending in M9 medium without carbon, and vortexing and triturating if necessary to homogenize. The individual cultures were then diluted to OD600 of 0.5 ± 0.05, combined at equal proportions, and inoculated in biological triplicate into the combinatorial media plates at final concentrations of OD600 0.05 in 300 μl. Each plate additionally contained three control wells: one uninoculated well with 50 mM C of D-glucose to control for contamination, and two inoculated wells with no nutrient to assess the decay of the initial inocula. The communities were grown at 30°C with shaking at 300 rpm for periods of 24 or 48 hours before each passage. At each passaging step, the cultures were triturated 10 times to ensure the communities were homogenized and 10 μl were transferred to 290 μl of fresh media for the subsequent growth period. Yields of the cultures were quantified by transferring 150 μl to clear 96-well plates (Corning, Corning, NY) and reading absorbance (OD600) using a microplate reader (BioTek-Synergy HTX). Biomass quantities are reported as the difference between the raw OD600 readings of each sample and the mean OD600 value of the negative control wells. Outlying OD600 readings were removed by calculating Z-scores *M* for each individual measurement *x*_*i*_ using the median absolute deviation (MAD):

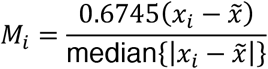

where 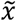 is the median across three biological replicates and 0.6745 represents the upper quartile of the normal standard distribution, to which the MAD converges. If the Z-score of an individual measurement exceeded 3.5, it was considered an outlier and removed. This process resulted in the elimination of 8 data points (out of 192) for com3, 15 for com4, 10 for com13, and 17 (out of 768) across the four monocultures. A summary of the organisms, nutrients, and culturing conditions for each experiment is found in Supplementary Table 5.

Communities to be sequenced were centrifuged at 4000 rpm for 10 minutes and the supernatant was removed. Cell pellets were stored at −20°C until DNA collection was performed using a 96-well genomic DNA purification kit (Invitrogen). To harvest the DNA, each cell pellet was resuspended in 180μl lysis buffer containing 25 mM Tris-HCl, 2.5 mM EDTA, 1% Triton X-100, and 20 mg/ml Lysozyme (Sigma-Aldrich). The samples were mixed by vortexing and incubated at 37°C for 30 minutes, after which 20mg/ml of RNase A (Invitrogen, Carlsbad, CA) and 20 mg/ml of Proteinase K (Invitrogen) with PureLink Pro 96 Genomic Lysis/Binding Buffer (Invitrogen) were added. The samples were mixed by vortexing and centrifuged after each reagent was added. The samples were incubated at 55°C for 30 minutes, after which 200 μl of 100% ethanol (Sigma-Aldrich) were added. DNA from the resulting lysates was purified using a vacuum manifold according to the purification kit protocol (Invitrogen). Purified DNA was normalized to 15 ng/μl using a NanoDrop 1000 spectrophotometer (Thermo Scientific, Waltham, MA). Library preparation was performed based on a paired-end approach developed by Preheim *et al*.^39^, which targets the V4 region of the 16s rRNA gene with the primers U515F (5’- GTGCCAGCMGCCGCGGTAA) and E786R (5’-GGACTACHVGGGTWTCTAAT). Libraries were sequenced using an Illumina MiSeq platform at either the MIT BioMicroCenter, Cambridge, MA (com13a) or at QuintaraBio, Boston, MA (com13). QIIME2^40^ was used to demultiplex raw files and produce FASTQ files for forward and reverse reads for each sample. DADA2^41^ was used to infer sequence variants from each sample and a naïve Bayes classifier was used to assign taxonomic identities using a custom reference database with a 95% confidence cutoff.

Communities to be assayed by agar plating were diluted by a factor of 10^4^ and spread on LB agar plates using autoclaved glass beads. Plates were prepared by autoclaving and distributing 18 mL of LB agar (Sigma-Aldrich) into petri dishes using a peristaltic pump (TriTech, Los Angeles, CA). Inoculated plates were incubated at 30°C and imaged after 72 hours using a flatbed scanner for colony counting. Colony counts for com3 and com4 were adjusted based on a standard dilution of the community members at equal concentrations measured by OD.

Significance between growth yields under differing environments was determined using a one-sided two-sample t-test with significance cutoffs of 0.05, 0.01, and 0.001. Species richness (*S*) is defined as the number of different organisms detected in a particular environment. Shannon entropy (*H*) is defined as follows:

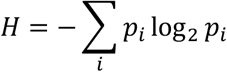

where *p*_*i*_ is the relative abundance of organism *i* in a sample.

For hierarchical clustering analysis of communities, Spearman correlation coefficients were computed either for pairs of environments or pairs of organisms based on normalized vectors of species abundances. Hierarchical clustering was performed on the correlation coefficients using the ‘clustergram’ function in MATLAB, which calculated distances between clusters using the UPGMA method based on Euclidean distance.

### Computation of nonlinearity scores for yield, species richness, and Shannon entropy

To quantify nonlinearities in how taxonomic diversity and balance could change in incrementally more complex environments, we first established definitions of expected values of species richness *S* and Shannon entropy *H* based on their values in lower-complexity conditions. Let a combined set of nutrients *AB* be defined as the union of nutrient sets *A* and *B*. For nutrient sets *A*, *B*, and *AB*, the vectors of species abundances in each set are defined as *V*_*A*_, *V*_*B*_, and *V*_*AB*_, respectively. The species richness values *S* for each set are therefore simply the number of positive species abundance values in each vector. Based on the organisms that survived in sets *A* and *B*, we establish the naïve assumption that at least as many organisms as survived in either environment will also survive in set *AB*. We therefore define *V*_*expected,AB*_ as max(*V*_*A*_, *V*_*B*_), and *S*_*expected,AB*_ as the number of organisms contained in this set. Our epistasis score for species richness *E*_*S,AB*_ becomes:

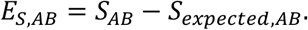

We use a similar expectation to calculate Shannon entropy epistasis *E*_*H*_, where . We first calculate the observed *H* for nutrient set *AB* as *H*_*AB*_ = − ∑ *V*_*AB*_ log_2_ *V*_*AB*_, and the expected *H* for nutrient set *AB* based on *A* and *B* as *H*_*expected,AB*_ = max (*H*_*A*_, *H*_*B*_). The epistasis score for Shannon entropy *E*_*H,AB*_ is therefore:

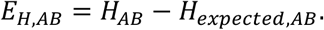

### Flux balance modeling

To estimate the number of secreted and absorbed metabolites in com4, we used experimentally-validated genome-scale models for each of the four organisms^42–45^. Genome-scale models are mathematical representations of the known metabolic capacities of individual organisms^46^. They are constrained by known maximum metabolic fluxes *v*_*max*_ through internal and transport reactions, as well as by reaction stoichiometric constraints represented by a matrix *S*. Flux balance analysis (FBA), a mathematical optimization technique, can then be applied to the models in order to define the metabolic fluxes *v* within the organism’s network that will maximize a particular objective, such as growth^47^. This technique allows us to interrogate the growth rate of organisms under specific environmental conditions, as well as rates of nutrient consumption and metabolite secretion.

Our application of FBA uses the COBRA toolbox^48^ and is largely based off of an implementation used in a previous study^20^. Here, we employed FBA to simulate the growth of the four organisms in com4 in the 63 combinatorial medium conditions we tested experimentally. We first defined an *in silico* M9 minimal medium consisting of the various inorganic molecules present in the *in vitro* minimal medium (Supplementary Table 7). These molecules were provided to the genome-scale models at nonlimiting availabilities by setting the corresponding maximum flux bounds *v*_*max*_ to 1000 mmol/gDW/hr. Depending on the environmental condition, we supplied each *in silico* organism with the appropriate carbon sources by setting the corresponding maximum flux bounds *v*_*max*_ to 10 mmol/gDW/hr. We then applied FBA by maximizing the growth rate and minimizing the sum of the fluxes in the network. This latter step was employed in order to more closely model proteome usage and minimize metabolite cycling throughout the network^49^. The optimization problem applied is therefore:

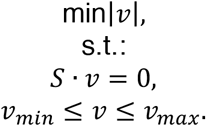

If an *in silico* organism grew on a given environmental condition, we recorded which organic metabolites were predicted to be taken up and secreted. These are summarized for all environments in Supplementary Table 8.

### Consumer resource modeling

We employed a dynamical modeling framework to simulate the yields of arbitrary communities in increasingly complex environments, as well as the relative abundances of com4. The model is an adaptation of Robert MacArthur’s consumer resource model^50–52^ and a subsequent modification by Marsland *et al.*^18^, which simulates the abundances of organisms over time as a function of resource availability, metabolic preferences, and exchange of secreted metabolites.

We define the individual species abundances as *N*_*i*_ for *i* = 1, …, *S*, and the resource abundances as *R*_*α*_ for *α* = 1, … *M*. The key variable in calculating the abundances *N*_*i*_ is a stoichiometric resource utilization matrix *C*_*iα*_, which defines the uptake rate per unit concentration of resource *α* by organism *i* (Figure 1e). To calculate the growth of each organism on each resource, we multiply this matrix by a Monod consumption term *R*_*α*_/ (*k*_*i,a*_ − *R*_*α*_) that simulates concentration-dependent resource depletion. Each consumed resource type *α* with abundance *R*_*α*_ is therefore consumed by organism *i* at a rate *C*_*iα*_ *R*_*α*_/ (*k*_*i,a*_ − *R*_*α*_). These resources *α* are then transformed into other resources *β* by the organisms via a species-specific normalized stoichiometric matrix *D*_*αβi*_. A fraction *l* of the resultant metabolic flux is returned to the environment to be made available to other organisms, while the rest is used for growth. In addition to these resource consumption terms, the species abundances are also defined by (i) a species-specific conversion factor from energy uptake to growth rate *g*_*i*_, (ii) a scaling term representing the energy content of each resource *w*_*α*_, (iii) a quantity representing the minimal energy uptake required for maintenance of each species *m*_*i*_, and (iv) a dilution rate *d*. These terms are further defined in Supplementary Table 7. Taken together, the species abundances *N*_*i*_ over time are defined by:

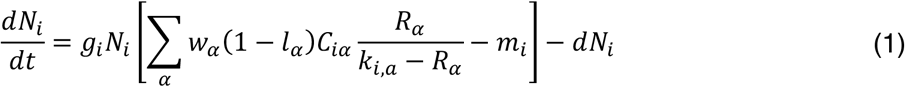

The resource abundances *R*_*α*_ are first defined by a continuous external supply rate *k*_*α*_ in addition to the constant dilution rate *d* (Supplementary Table 9). Each resource is then consumed, in a manner dependent on the matrix *C*_*iα*_, and converted to other nutrients based on the stoichiometric matrix *D*_*αβi*_:

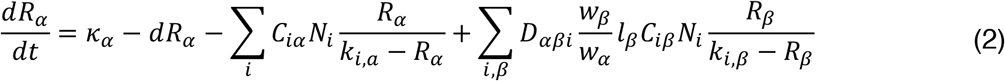

We selected the parameters for our equations based on experimental observations and quantities obtained from the literature. The dilution rate *d* was based on the 48-hour dilution timeframe we used in our experiments, in which 10 μl of culture was passaged into a total of 300 μl of fresh, uninoculated media. The resource input rate *k*_*α*_ was defined similarly for all nutrients, equaling the initial concentration of each nutrient divided by a 48-hour dilution timeframe. Kinetic growth curves for com14 (Supplementary Table 5, Supplementary Figure 17a) were used to estimate the orders of magnitude for the remaining parameters, based on the community reaching an average of approximately 2.4 × 10^8^ CFU/mL (OD600 0.3) within 20 hours in 50 mM C of D-glucose. The conversion factor from energy uptake to growth rate *g*_*i*_, as well as the energy content of each resource *w*_*α*_, were set to 1 and 1 × 10^8^, respectively, in order to approximate this magnitude of growth. The Monod resource consumption half-velocity constant was set to 3000 g/mL for all nutrients in order to approximate the experimentally-observed growth timeframe. Lastly, the minimum energy requirements for all organisms *m*_*i*_ were informed by the community yields at steady state and the leakage fraction *l*_*α*_ was set to 0.8 based on community simulations in Marsland *et al.*^18^. These quantities are summarized in Supplementary Table 9.

In our initial CRM simulations, in which we generated a null model for community yield under increasing environmental complexity (Supplementary Figure 5), we simulated the growth of a community containing *S* = 13 arbitrary organisms. These organisms were defined by randomly-populated nutrient preference matrices *C*, in which each organism had a 50% chance of being able to utilize a particular nutrient. In this way, the organisms had comparatively even, though not necessarily intersecting, nutrient utilization profiles. Each community was simulated 50 times in each environment, so that the nutrient preference matrices could be randomly repopulated. This process allowed us to more effectively sample the large space of possible nutrient utilization matrices and obtain a clearer indication of how mean community yields changed in response to increasing environmental complexity. The environments were generated from a set of *M* = 32 arbitrary nutrients, which were combined in a scheme similar to that of com3, com4, and com13: 32 conditions with one nutrient, 16 conditions with two nutrients, and so on, up until one condition with all 32 nutrients. To test the effects of metabolic exchange, our simulations contained between 1 and 10 unique secreted metabolites, which could also be consumed by organisms according to randomly-defined preferences in *C*. For a more realistic representation of metabolic conversion, these byproducts were matched with primary nutrients in conversion matrix *D*, which was randomly populated according to a transition probability of 0.25, meaning that a given metabolic byproduct had a 25% chance of being converted from a given primary nutrient. This matrix was normalized across each primary nutrient to ensure conservation of mass. Our results for yield, species richness, and Shannon entropy are presented as the average across all quantities of metabolic byproducts. The initial species and resource abundances were set to 6 × 10^6^ CFU/mL and 1.5 g/mL, respectively, to approximate the initial OD600 of 0.05 and the initial nutrient concentration of 50 mM C of glucose used in our experiments. We then simulated the growth of these communities over the course of 288 hours (based on com3, com4, and com13 culturing timescale) with a timestep of 0.01 hours and quantified their yields at the endpoints.

For our simulations of species dynamics in com3 and com4, we parametrized the nutrient preference matrix *C* based on the growth yields of each individual organism on the 32 individual carbon sources (Supplementary Figure 7b). To determine the fraction of each nutrient *α* that could be converted to a secreted metabolite *β*, we used flux-balance models for each of the four relevant organisms^42–45^. We calculated the secretion fluxes of metabolic byproducts from the organisms under each of the 32 individual nutrients using the technique described above, and used the ratio of secretion to intake fluxes to populate the *D* matrix. This matrix was then normalized across each primary nutrient to ensure conservation of mass. Nutrient uptake efficiencies for these secreted metabolites were then assigned according to the same monoculture growth data used to define preferences for the primary nutrients. As with our initial arbitrary communities, initial species abundances were set to 6 × 10^6^ CFU/mL and initial nutrient abundances were set to 1.5 g/mL. We then simulated the growth and potential metabolic exchange of all four community members in coculture over the course of 288 hours with a timestep of 0.01 hours in the 63 environmental conditions for com3 and com4 (Supplementary Table 3).

To simulate the growth of randomly-defined communities containing generalist and specialist organisms, we again relied on random sampling to populate the resource preference matrix *C*. For our 4-species communities, we used a more stringent generalist-specialist designation based on how many nutrients were able to be consumed by the top organisms in our Biolog assay: an organism was classed as a generalist if it was able to grow on more than 90% of the nutrients (e.g. *P. aeruginosa)*, and a specialist if it was able to grow on fewer than 50% of the nutrients (e.g. *B. subtilis*). Therefore, if an organism was a generalist it had a 90% chance of consuming any given nutrient, versus a 50% chance if it was a specialist. For our first 13-species community, each organism had a probability of 0.5 of utilizing a particular nutrient. In our second 13-species community, this probability was defined by the proportion of nutrients each com13 organism was able to utilize in our Biolog assay. For the nutrient conversion matrix *D*, we established a fixed probability of a particular organism *i* converting a nutrient *α* into a metabolite *β* of 25%. We carried out 50 random simulations of these communities, randomly repopulating the *C* and *D* matrices each time to adequately sample the possible space of nutrient preferences and conversions. These simulations were also run with the same parameters, initial conditions, and timeframe as those for com4.

## Acknowledgements

We thank Melisa L. Osborne, Christopher P. Mancuso, Jennifer M. Bhatnagar, Sylvie Estrela, and Ilija Dukovski for experimental advice and technical assistance. We are also grateful to Sean P. Mullen for access to the robotic liquid-handling system and to James E. Fifer for relevant experimental training. Additionally, we thank past and present members of the Segrè group for helpful discussions on the research and the manuscript, particularly David B. Bernstein, Joshua E. Goldford, Robert Marsland III, Demetrius DiMucci, Elena Forchielli, and Devlin Moyer. A.R.P. is supported by a Howard Hughes Medical Institute Gilliam Fellowship and a National Academies of Sciences, Engineering, and Medicine Ford Foundation Predoctoral Fellowship. We gratefully acknowledge support from the U.S. Department of Energy, Office of Science, Office of Biological & Environmental Research through the Microbial Community Analysis and Functional Evaluation in Soils SFA Program (m-CAFEs) under contract number DE-AC02-05CH11231 to Lawrence Berkeley National Laboratory, as well as the National Institutes of Health (grants 5R01DE024468, R01GM121950), the National Science Foundation (grants 1457695 and NSFOCE-BSF 1635070), the Human Frontiers Science Program (grant RGP0020/2016), and the Boston University Interdisciplinary Biomedical Research Office.

## Author contributions

A.R.P and D.S. designed the research. A.R.P. designed and performed experiments, collected data, wrote data analysis code, developed the models, and ran and analyzed simulations. A.R.P. and D.S. wrote the manuscript. Both authors read and approved the final manuscript.

## Declaration of interests

The authors declare that no competing interests exist in relation to this manuscript.

## Supplementary Information

### Supplementary Text

#### Drivers of yield epistasis in communities of varying sizes

Our observation that the yields of individual organisms and small communities increased with environmental complexity – but those of larger communities remained constant – suggests an interplay between different mechanisms that either enhance or dampen *E*_*Y*_ based on initial community size. On the one hand, community experiments (Supplementary Figure 17) and previous studies^14,20^ have indicated that microbial growth efficiency can scale nonlinearly with concentration, and that community growth rates can increase with environmental complexity (Supplementary Figure 18a)^3,53^. Such metabolic nonlinearities may be more dominant in smaller communities given that they can be more commonly dominated by a single organism (Supplementary Figure 23). On the other hand, ecological phenomena such as cross-feeding could make a wider pool of nutrients available over time, enriching even simple environments^12^ and thereby reducing the positive impact that initially complex environments have on yields. Given that they contain a greater variety of organisms, it may be that larger communities allow for the accumulation of more of these nutrients, potentially explaining the lack of skewness in *E*_*Y*_. This notion is supported by our observation that the yields of a different 13-species community grown in fewer carbon sources (com13a) did not significantly increase with environmental complexity (Supplementary Figure 18b), as well as by the distribution of *E*_*Y*_ for com13 being skewed at earlier experimental timepoints (Supplementary Figure 19). In addition, we observed evidence of possible byproduct utilization in the form of diauxic shifts in batch culture experiments (Supplementary Figure 3a, Supplementary Figure 17a-c). Nonetheless, it is not clear whether the number of secreted metabolic byproducts would be expected to increase significantly with environmental complexity. In fact, stoichiometric modeling suggested that the number of secreted metabolites quickly plateaus as the number of resources increases Supplementary Figure 20). Furthermore, such an amplification of the space of available nutrients would result in increased taxonomic diversity^20,54^, which was not observed experimentally.

#### Generating specific environment-phenotype mappings

Given the taxonomic variability observed across our dataset, we suspected that unrelated nutrient combinations could nonintuitively yield similar taxonomic compositions. A hierarchical clustering of all 63 nutrient combinations revealed such environment-phenotype pairings (Supplementary Figure 21), which also resulted in the emergence of three distinct organism groupings (Supplementary Figure 22). These groupings, which resemble previously-identified family-level functional relationships in natural communities^12^, provide insight into the types of nutrients that need to be present to favor a particular taxon. They do not, however, explain how individual carbon sources behave in higher-order combinations. We therefore generated a linear model to determine whether any particular carbon sources were more universally associated with higher taxonomic diversity. Indeed, although we identified a number of such nutrients, we still found that their effects could be eclipsed by those that disproportionately favored a single organism (Supplementary Table 6).

#### Modeling the effects of generalists on community diversity

While individual nutrient combinations can explain specific community properties, the lack of an observed relationship between environmental complexity and taxonomic diversity prompted us to hypothesize whether the patterns we identified could be explained by a broader ecological principle. Specifically, we sought to determine whether some properties of the nutrient use capabilities of our organisms could account for the low degrees of taxonomic diversity we observed. We therefore first used our consumer resource model to simulate the taxonomic compositions of com3 and com4, which were strikingly low in biodiversity and often dominated by the organisms with the broadest nutrient utilization capabilities (Supplementary Figure 23). Our model was parametrized with experimentally-obtained growth data (Supplementary Figure 7b) and featured the potential for cross-feeding of secreted byproducts^12,18^ informed by flux-balance predictions of metabolic turnover (Supplementary Figure 20a). This parametrization enabled us to make quantitative estimates of community growth trajectories and metabolic exchange (Supplementary Figure 24), yielding accurate predictions of the dominance of *P. aeruginosa* – which had the broadest set of usable carbon sources – across most conditions in com4. However, our model could not fully explain the dominance of *S. oneidensis* in com3 or the increases in yield we observed experimentally (Supplementary Figure 25).

Despite some inaccuracies, our experimentally-parametrized model recapitulated the low levels of taxonomic diversity we observed *in vitro*. We thus asked whether, beyond the specific details of our small consortia, this effect could be explained by the resource utilization capabilities of a community’s constituent organisms, as previously suggested based on field observations^17^. To do this, we generated simulated 4-species communities whose constituent organisms had either uniform or uneven nutrient use capabilities determined via random sampling. These models showed that the presence of a generalist organism robustly decreased community diversity, in a way that strongly depended on the presence (but not necessarily quantity) of available secreted metabolites (Supplementary Figure 15a-d).

### Supplementary Figures

**Supplementary Figure 1.**
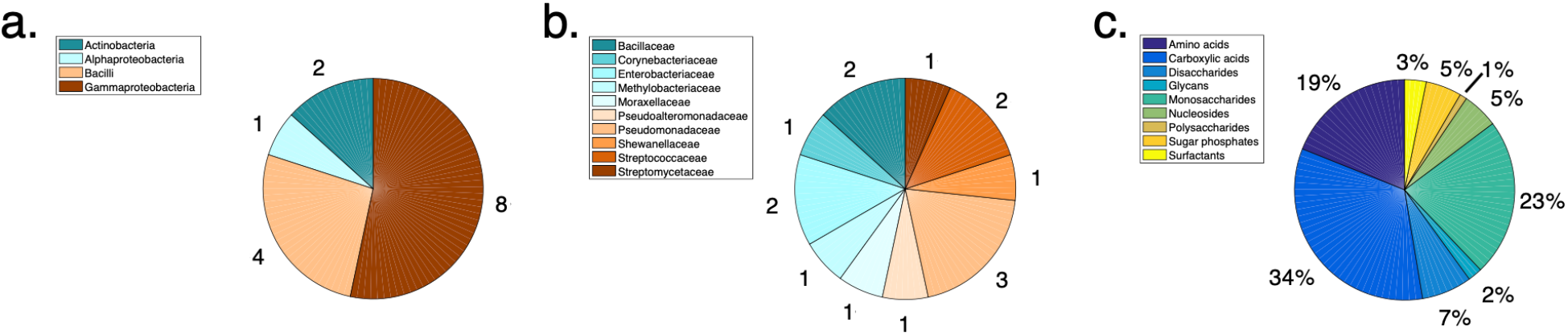
Organisms and carbon sources used in experiments. **(a-b).** Phylum- **(a)** and family-level **(b)** groupings of 15 bacterial organisms. **(c).** Nutrient categories for 95 carbon sources.

**Supplementary Figure 2.**
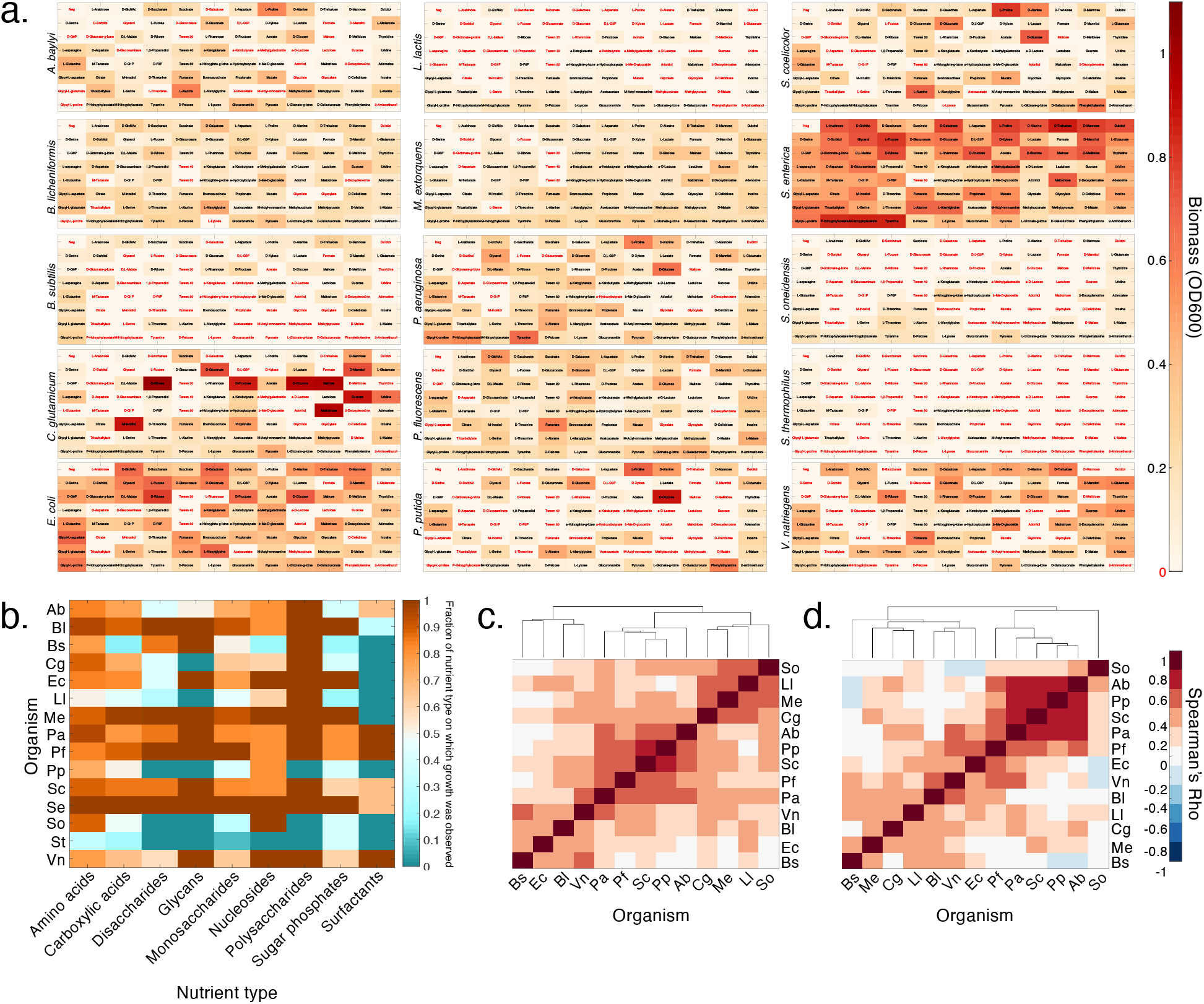
Results of Biolog phenotypic assay. **(a).** Average (3-replicate) species-specific growth profiles on Biolog carbon sources in M9 minimal medium after 48 hours. Raw OD600 values were corrected for liquid evaporation and a significance cutoff was applied to determine growth above the levels of the negative controls. Nutrients on which growth was not observed are marked in red. **(b).** Growth capabilities of all 15 organisms on 95 Biolog nutrients. Values displayed are the fraction of a specific nutrient type on which an organism displayed growth. **(c, d).** Hierarchical clustering of 13 selected organisms based on Spearman correlations of growth profiles on all 95 **(c)** and on 32 selected nutrients **(d)**. Organisms are abbreviated as: Ab: *A. baylyi*, Bl: *B. licheniformis*, Bs: *B. subtilis*, Cg: *C. glutamicum*, Ec: *E. coli*, Ll: *L. lactis*, Me: *M. extorquens*, Pa: *P. aeruginosa*, Pf: *P. fluorescens*, Pp: *P. putida*, Sc: *S. coelicolor*, So: *S. oneidensis*, Vn: *V. natriegens*.

**Supplementary Figure 3.**
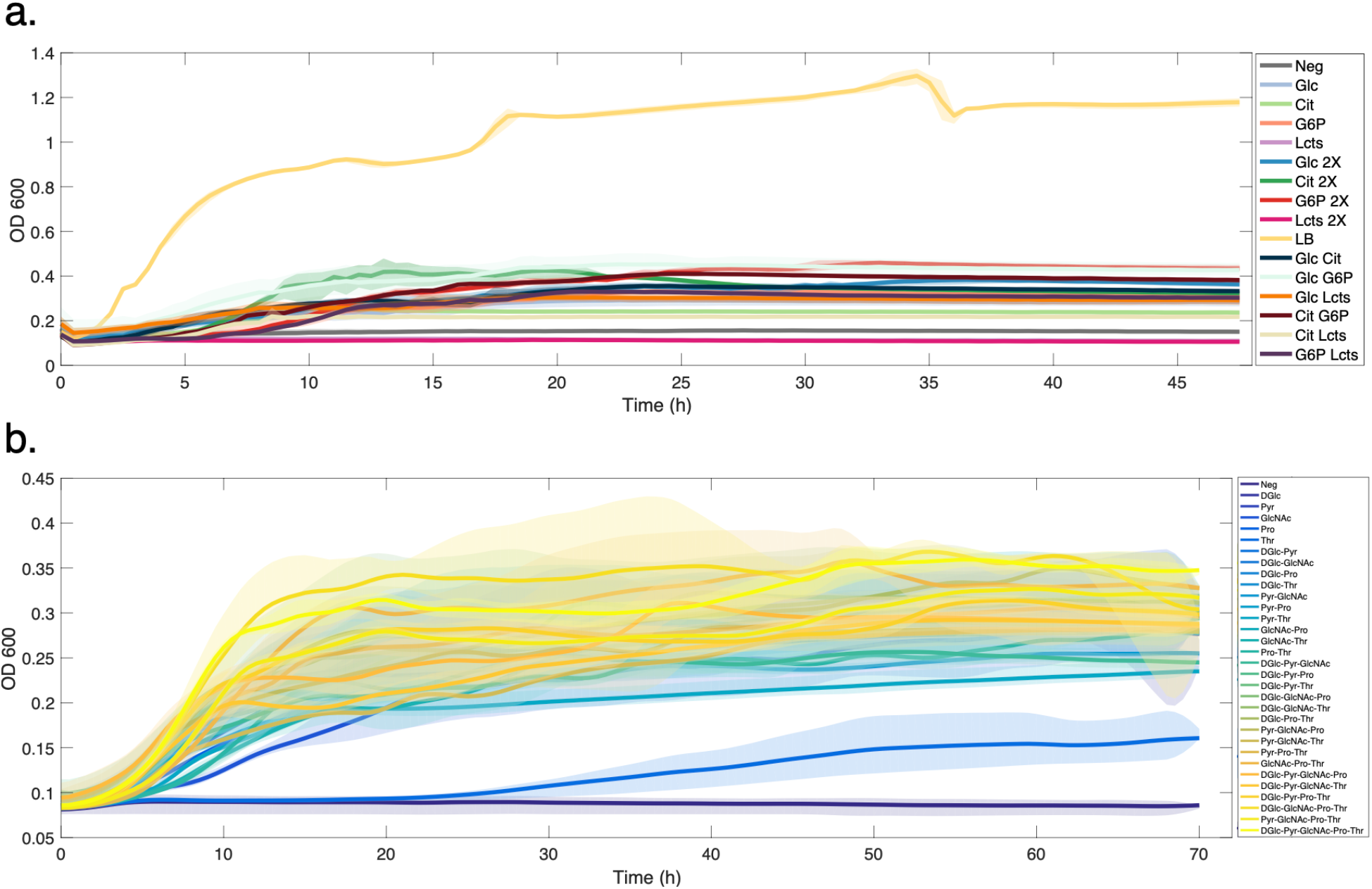
Growth curves for multispecies communities in combinatorial environments. **(a).** Growth trajectories of 14-species communities (com14, Supplementary Table 5) on combinations and different concentrations (25 mM C and 50 mM C) of D-glucose (Glc), citrate (Cit), glucose-6-phosphate (G6P), and a-D-lactose (Lcts). Double-carbon source conditions contain 50 mM C of total carbon source, for 25 mM C of each individual nutrient. **(b).** Growth trajectories of 13-species community (com13a) on equimolar concentrations (50 mM C) of five carbon sources: D-glucose (DGlc), pyruvate (Pyr), GlcNAc (GlcNAc), L-proline (Pro), and L-threonine (Thr). Shaded regions in both plots indicate standard deviations across three replicates per condition.

**Supplementary Figure 4.**
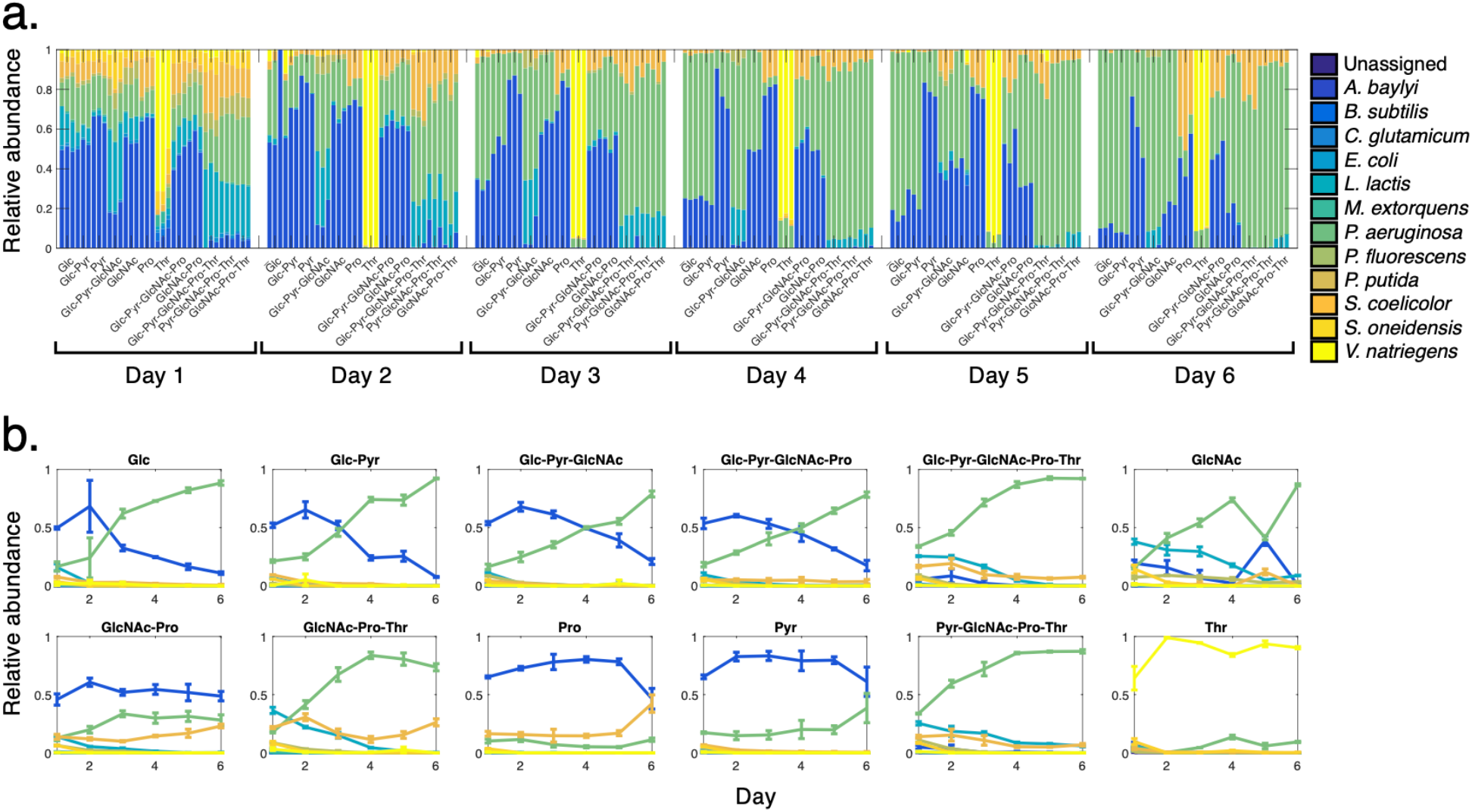
Taxonomic data for 13-species community grown on five carbon sources (com13a, Supplementary Table 5): D-glucose (DGlc), pyruvate (Pyr), GlcNAc (GlcNAc), L-proline (Pro), and L-threonine (Thr). **(a).** Relative abundance plots of all replicates over time. **(b).** Species relative abundance trajectories over time. Error bars indicate standard deviation.

**Supplementary Figure 5.**
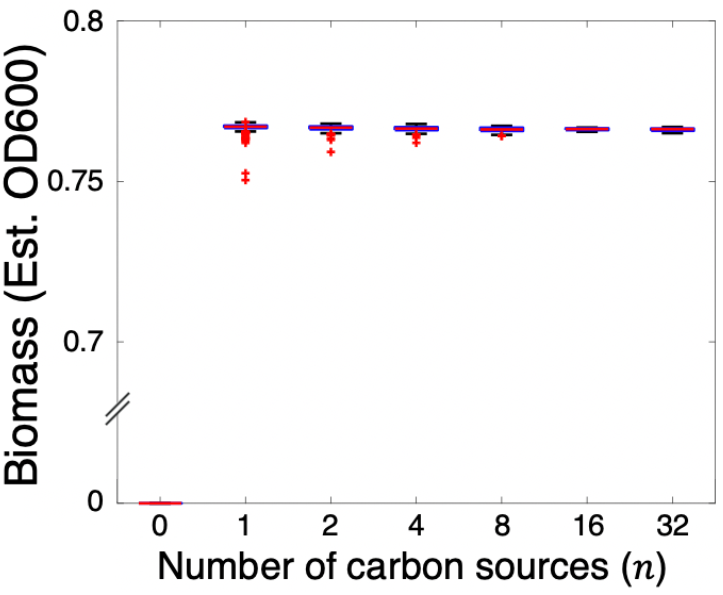
Consumer-resource model-predicted yields for simulated 13-member communities. Final biomass values after simulated 288 hours of community growth (corresponding to the full experimental timescale of com3, com4, and com13) on different combinations of nutrients. Here, final community yields converge to a median based on the total nutrient concentration (approximately 3 × 10^8^ CFU/mL, where an OD600 of 1 is estimated at 8 × 10^8^ CFU/mL). Here, as with all subsequent boxplots, the red central mark indicates the median, the top and bottom box edges indicate the 25^th^ and 75^th^ percentiles, respectively, the whiskers extend to the most extreme points not considered outliers, and the red ‘+’ symbols indicate outliers plotted individually. Paired t-tests showed that no significant changes in yield occurred with increasing environmental complexity (*p* = 0.07 for 1 vs. 2 nutrients, 0.08 for 2 vs. 4 nutrients, 0.13 for 4 vs. 8 nutrients, 0.71 for 8 vs. 16 nutrients, and 0.9 for 16 vs. 32 nutrients).

**Supplementary Figure 6.**
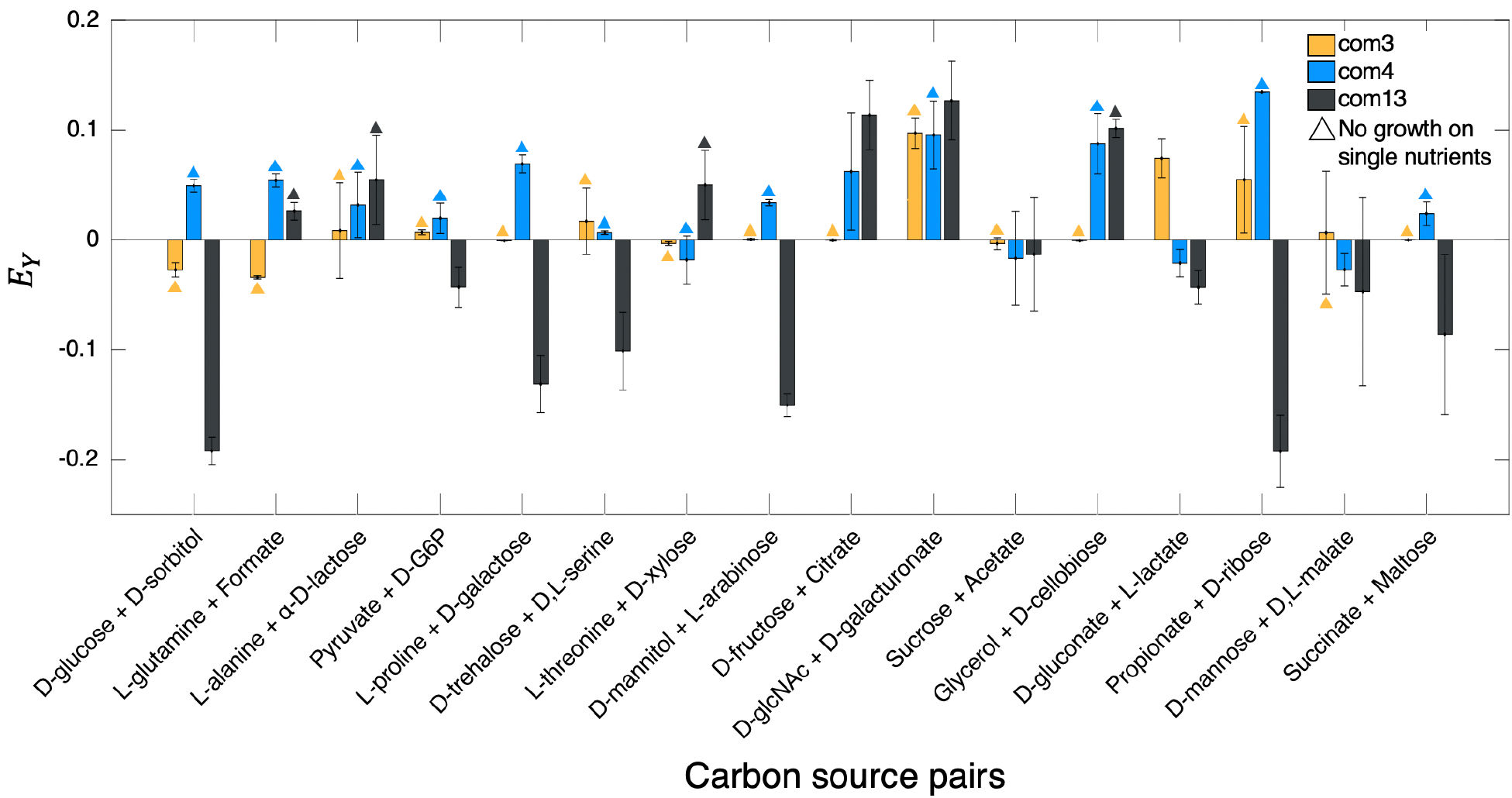
Yield epistasis *E*_*Y*_ for com3, com4, and com13 between community yields on pairs of carbon sources and yields on the corresponding single carbon sources. Error bars indicate standard deviation.

**Supplementary Figure 7.**
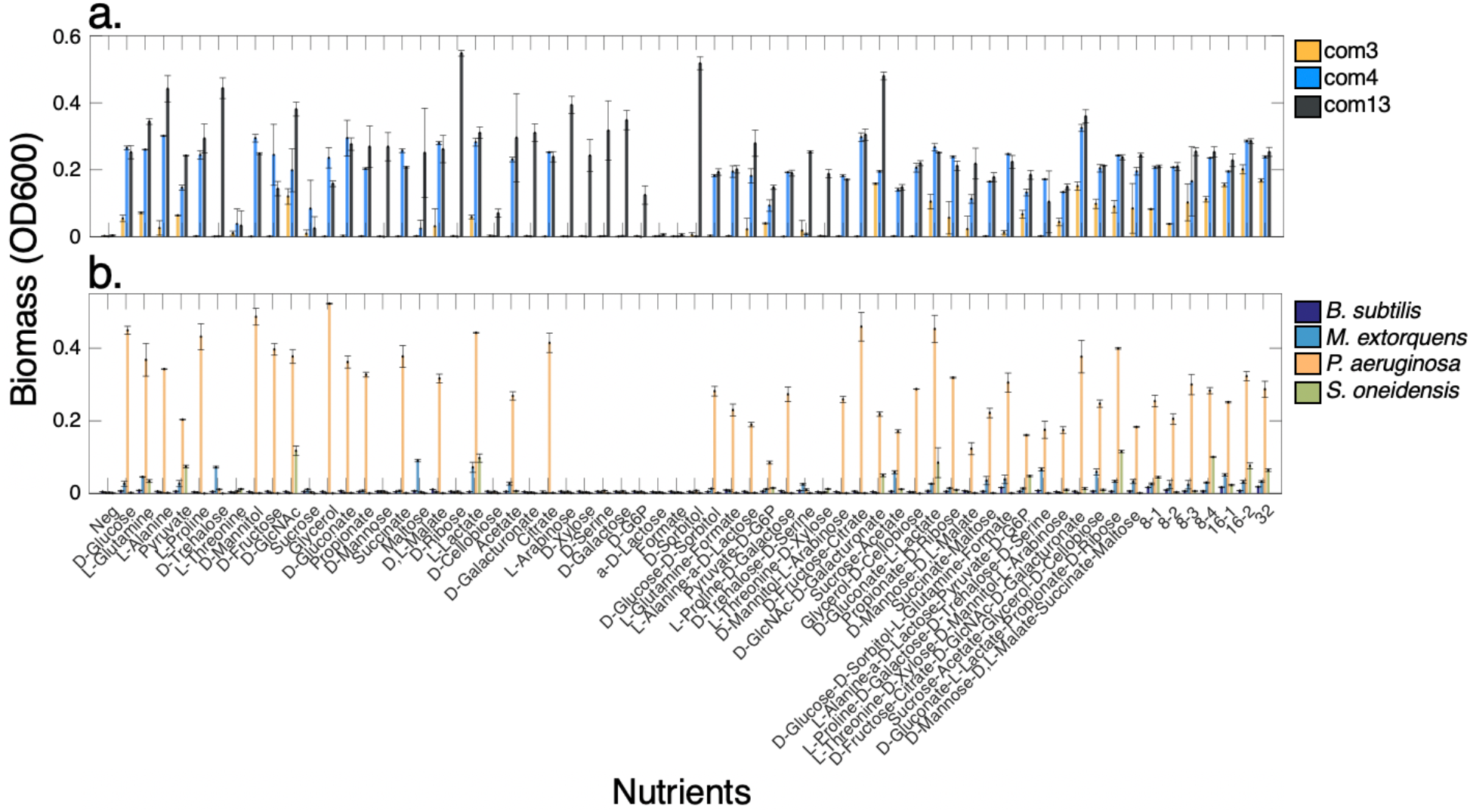
Endpoint nutrient combination-specific growth yields for multispecies communities com3, com4, and com13 **(a)** (Supplementary Table 5) and single organisms **(b)** on combinations of up to 32 carbon sources. Error bars indicate standard deviation.

**Supplementary Figure 8.**
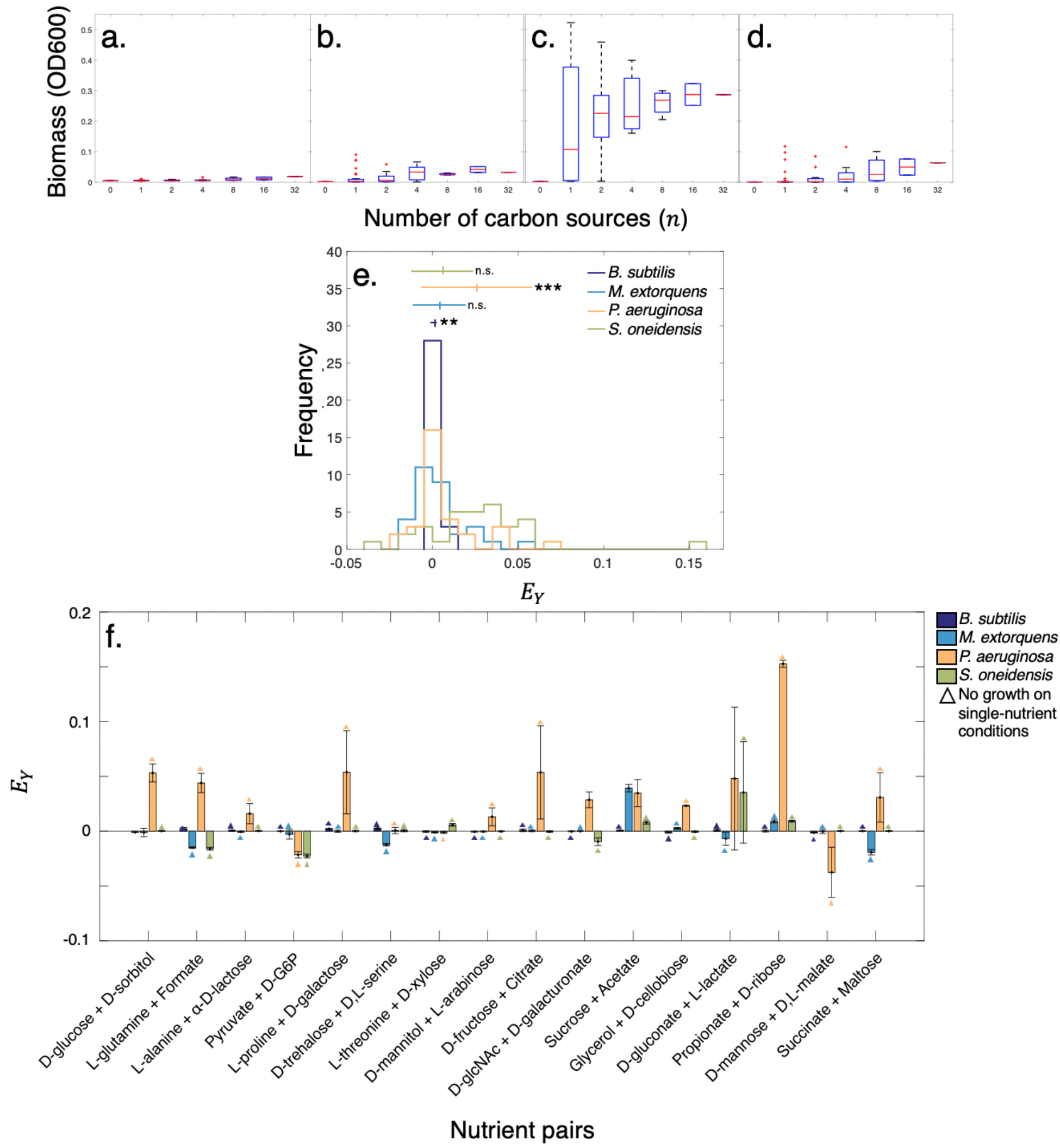
Monoculture biomass yields in combinatorial nutrients for **(a)** *B. subtilis*, **(b)** *M. extorquens*, **(c)** *P. aeruginosa*, and **(d)** *S. oneidensis*. **(e)**. Distributions of yield epistasis *E*_*Y*_ for four organisms. Bars and notches indicate mean and standard deviation. Significance determined using a one-sample t-test against a mean of zero and is indicated by (*) p < 0.05, (**) p < 0.01, and (***) p < 0.001. P-values for organisms are: 0.007 (*B. subtilis*), 0.122 (*M. extorquens*), 1.05e^−4^ (*P. aeruginosa*), and 0.056 (*S. oneidensis*). **(f).** Yield epistasis *E*_*Y*_ for four single organisms between pairs of carbon sources and corresponding single carbon sources. Error bars indicate standard deviation.

**Supplementary Figure 9.**
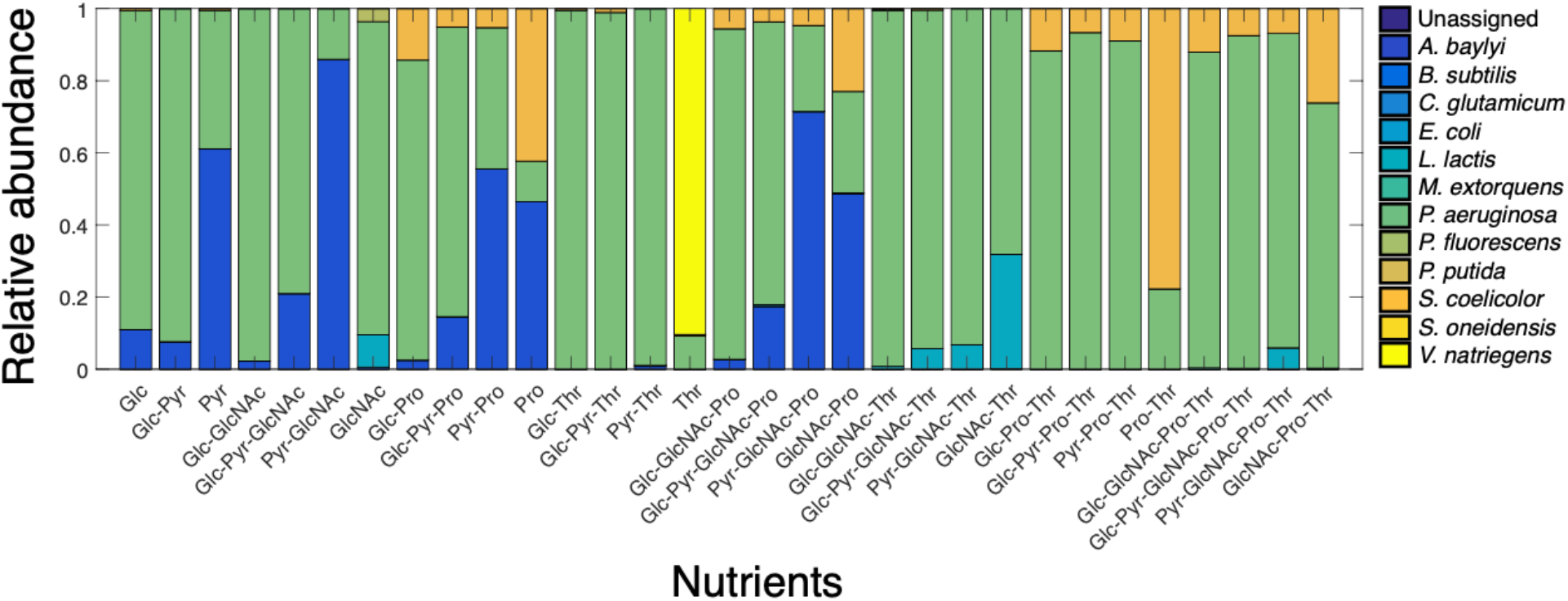
Mean relative abundances (averaged over 3 replicates) for 13-species community grown on five carbon sources (com13a): D-glucose (DGlc), pyruvate (Pyr), GlcNAc (GlcNAc), L-proline (Pro), and L-threonine (Thr).

**Supplementary Figure 10.**
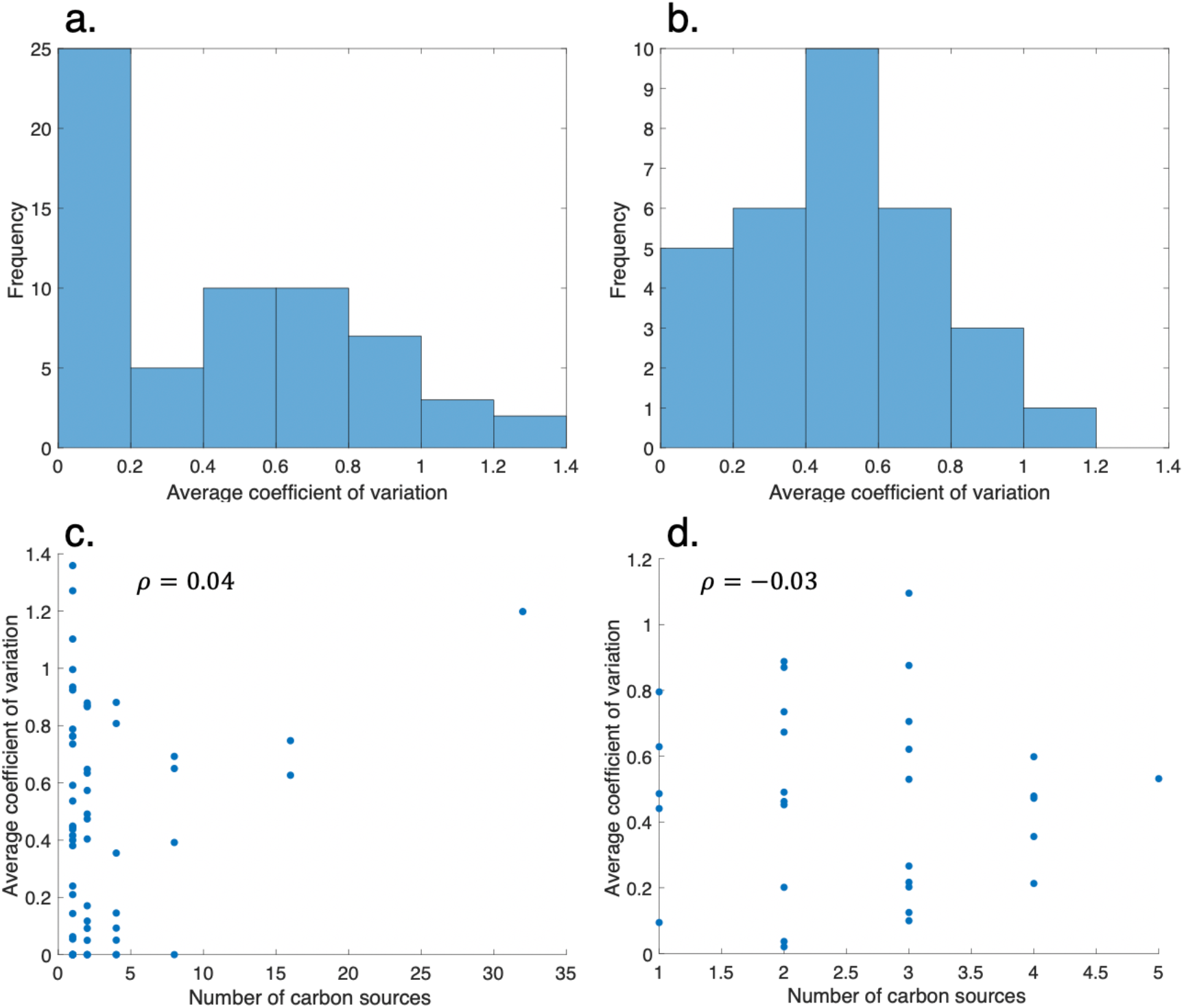
Distributions of inter-replicate coefficients of variation for 13-species community grown on 32 carbon sources (com13, **a**) and 13-species community grown on 5 carbon sources (com13a, **b**). **(c, d).** Inter-replicate coefficients of variation vs. number of carbon sources for com13 **(c)** and com13a **(d)** with Spearman correlation coefficients *ρ*. No significant correlations were found between inter-replicate variability and environmental complexity for either community (*p* = 0.73 for com13 and 0.86 for com13a).

**Supplementary Figure 11.**
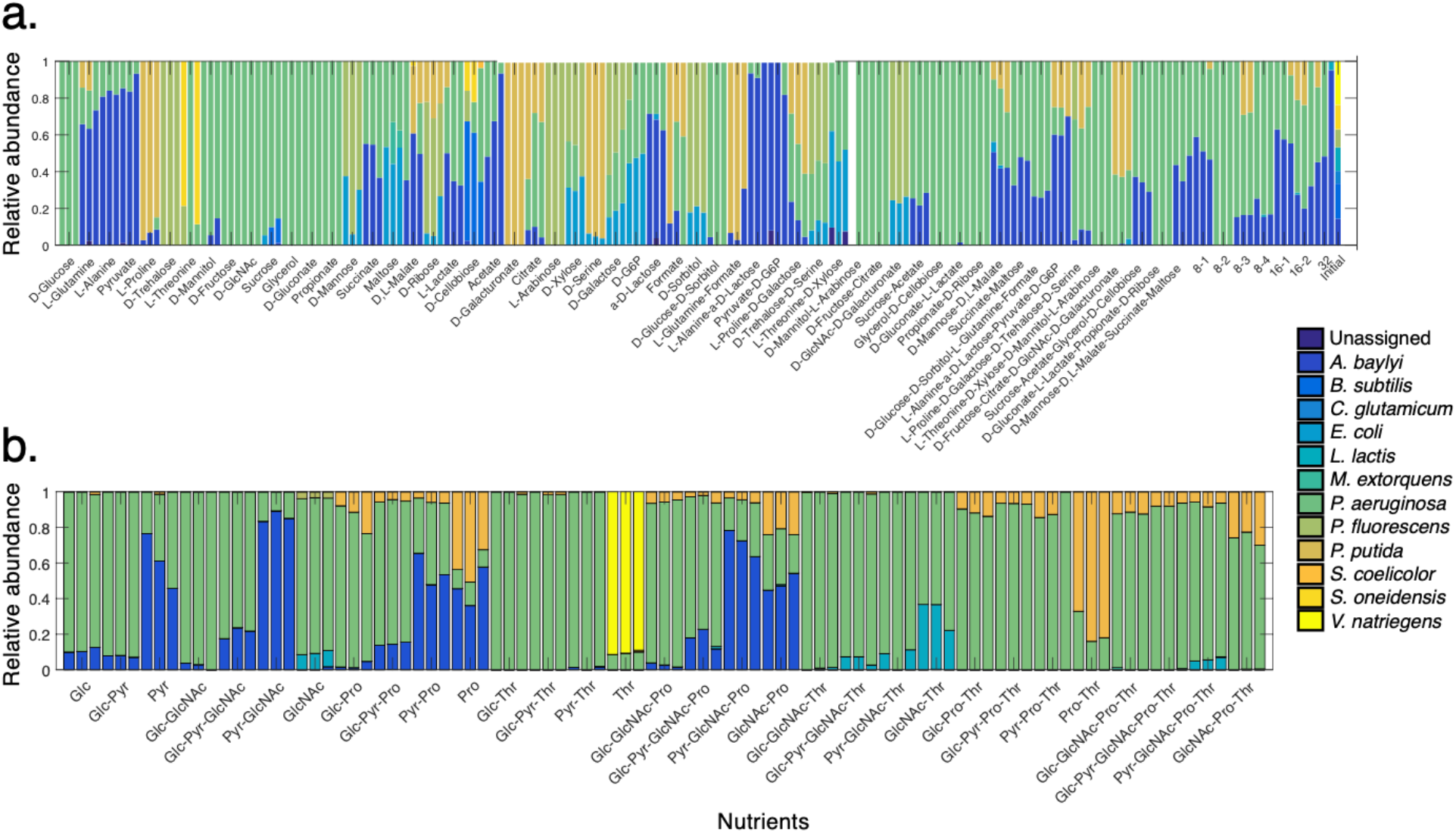
Endpoint taxonomic distributions over all replicates for 13-species community grown on 32 carbon sources (com13, **a**) and 13-species community grown on 5 carbon sources (com13a, **b**). We encountered general consistency in community composition between comparable conditions of the two 13-species experiments. The D-glucose, pyruvate, and D-GlcNAc conditions had the same dominant organisms in both experiments (*P. aeruginosa*, *A. baylyi*, and *P. aeruginosa*, respectively), and the L-proline condition was composed of *A. baylyi*, *P. aeruginosa*, and a third organism in both experiments with only the identity of the third organism being different (*P. putida* in com13 and *S. coelicolor* in com13a). Nonetheless, in com13, L-threonine resulted in a dominance of *S. oneidensis* in two of the replicates and of *P. aeruginosa* in one replicate, while com13a resulted in dominance of *V. natriegens* in all three replicates. However, com13 grew very minimally in L-threonine (OD600 0.03 ± 0.04) in comparison to com13a (0.15 ± 0.02), in addition to having a very high inter-replicate coefficient of variation for this condition (1.36), suggesting that it is an outlier.

**Supplementary Figure 12.**
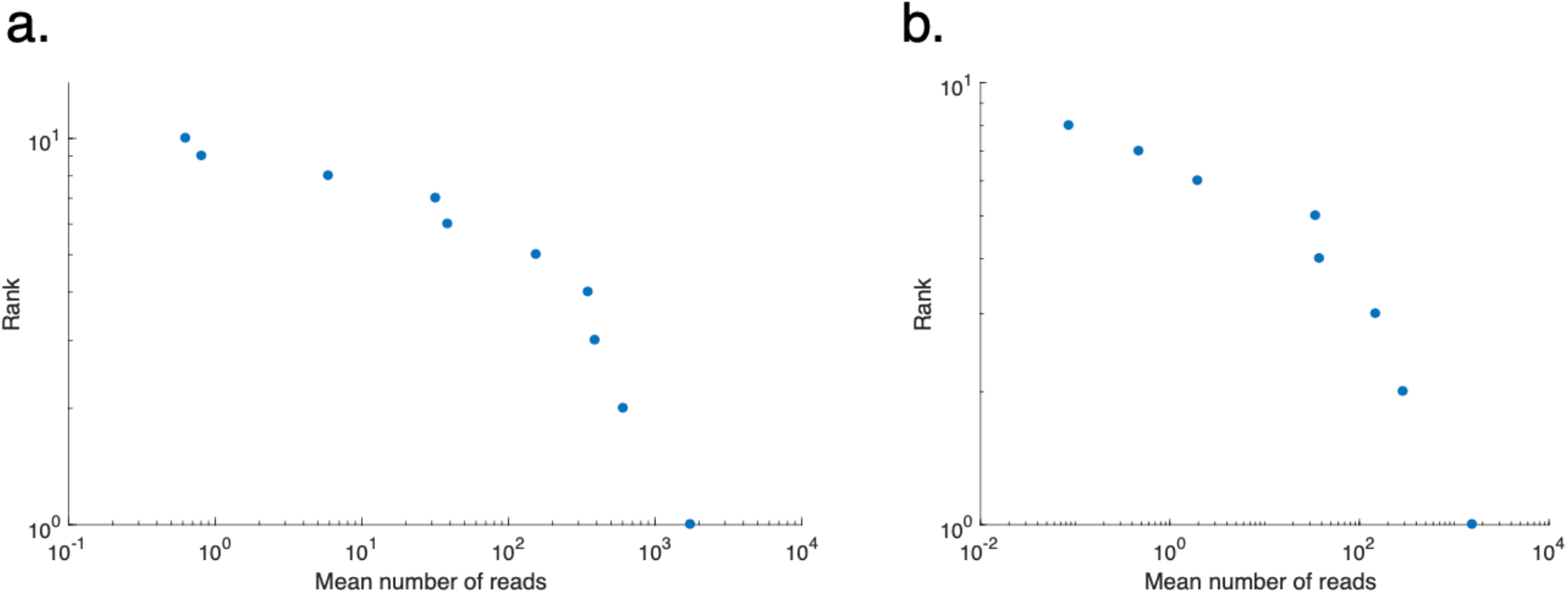
Rank-abundance plots for 13-species community grown on 32 carbon sources (com13, **a**) and 13-species community grown on 5 carbon sources (com13a, **b**). These relationships displayed decay patterns separated at characteristic scales of approximately 10^2^ reads, resembling double-power law relationships previously observed in a variety of natural ecosystems orders of magnitude more complex than our model communities ^22–24^. Despite difference in scale, this rank-abundance relationship suggests fundamental structural similarities in community composition across experimental systems. Moreover, this similarity extends to the scaling and prevalence of very low-abundance taxa, indicating the abundances of these community members are accurately represented within our populations.

**Supplementary Figure 13.**
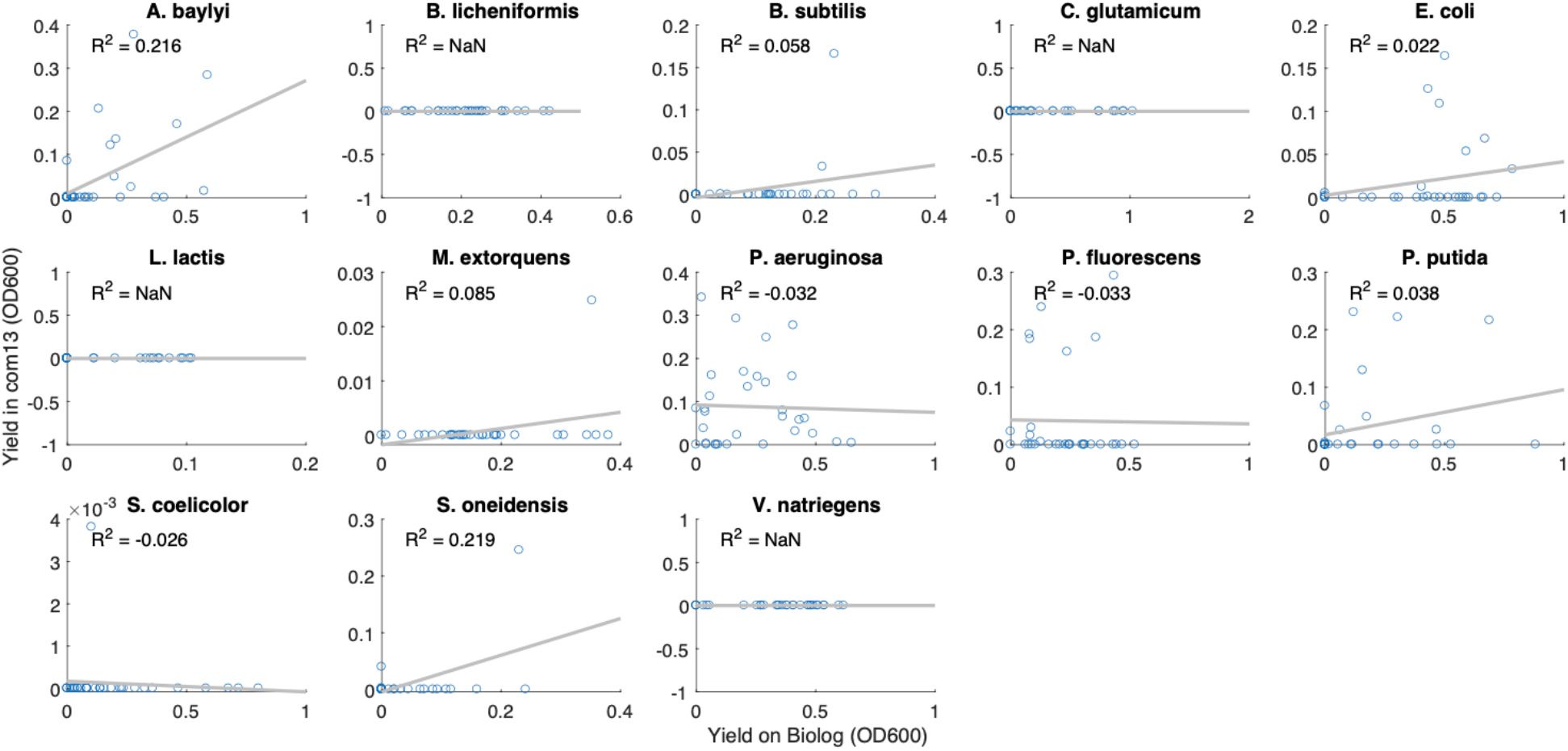
Organism-specific yields in 13-species community grown on 32 carbon sources (com13) vs. in monoculture, with best-fit lines calculated using ordinary least squares regression.

**Supplementary Figure 14.**
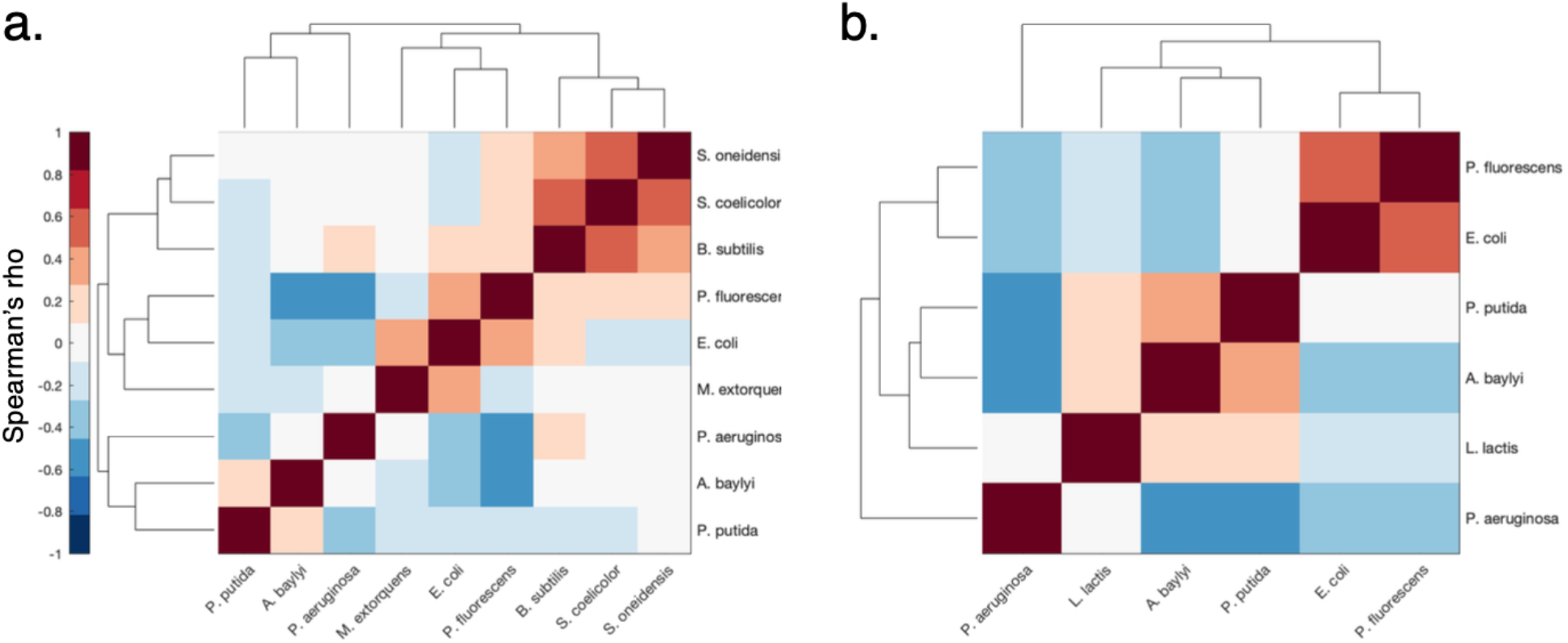
Hierarchical clustering of Spearman correlations between organism relative abundances across com13 environments (Supplementary Table 3). **(a).** Clustering of species-species correlations across single-nutrient conditions. We found that the overall structure of the species-species clusters in mixed cultures was dramatically different from that of the monocultures (Supplementary Figure 2d), with much lower degrees of interspecies similarities. In com13, higher degrees of similarity were observed between *B. subtilis*, *S. coelicolor*, and *S. oneidensis*, likely due to their co-occurrence in D-cellobiose. This similarity was not observed in the monoculture data, in which *B. subtilis* correlated more strongly with organisms that did not remain in com13. The species-species clustering of com13 also highlighted the profound dissimilarities between *P. fluorescens* and both *A baylyi* and *P. aeruginosa*. These anticorrelations contrast with the monoculture data, as *P. fluorescens* had displayed relatively high degrees of similarity with both organisms. This difference further clarifies the competitive effects observed between *P. fluorescens* and these two organisms in a community setting, which leads to exclusion of specific organisms despite their ability to utilize the provided nutrients. **(b).** Clustering of species-species correlations across multiple-nutrient conditions. We observed that the similarities between *E. coli* and *P. fluorescens* were more pronounced than in the single-nutrient conditions, reflecting the ability of these two organisms to coexist in across different environments (e.g. D-trehalose + D-serine and D-glcNAc + D-galacturonate). Despite also having the ability to coexist in more complex environments, the correlations between *A. baylyi* and *P. aeruginosa* decreased in the multiple-nutrient conditions. In fact, *P. aeruginosa* was found to be the most dissimilar organism in these environments, likely due to its ability to exclude all other organisms in many conditions.

**Supplementary Figure 15.**
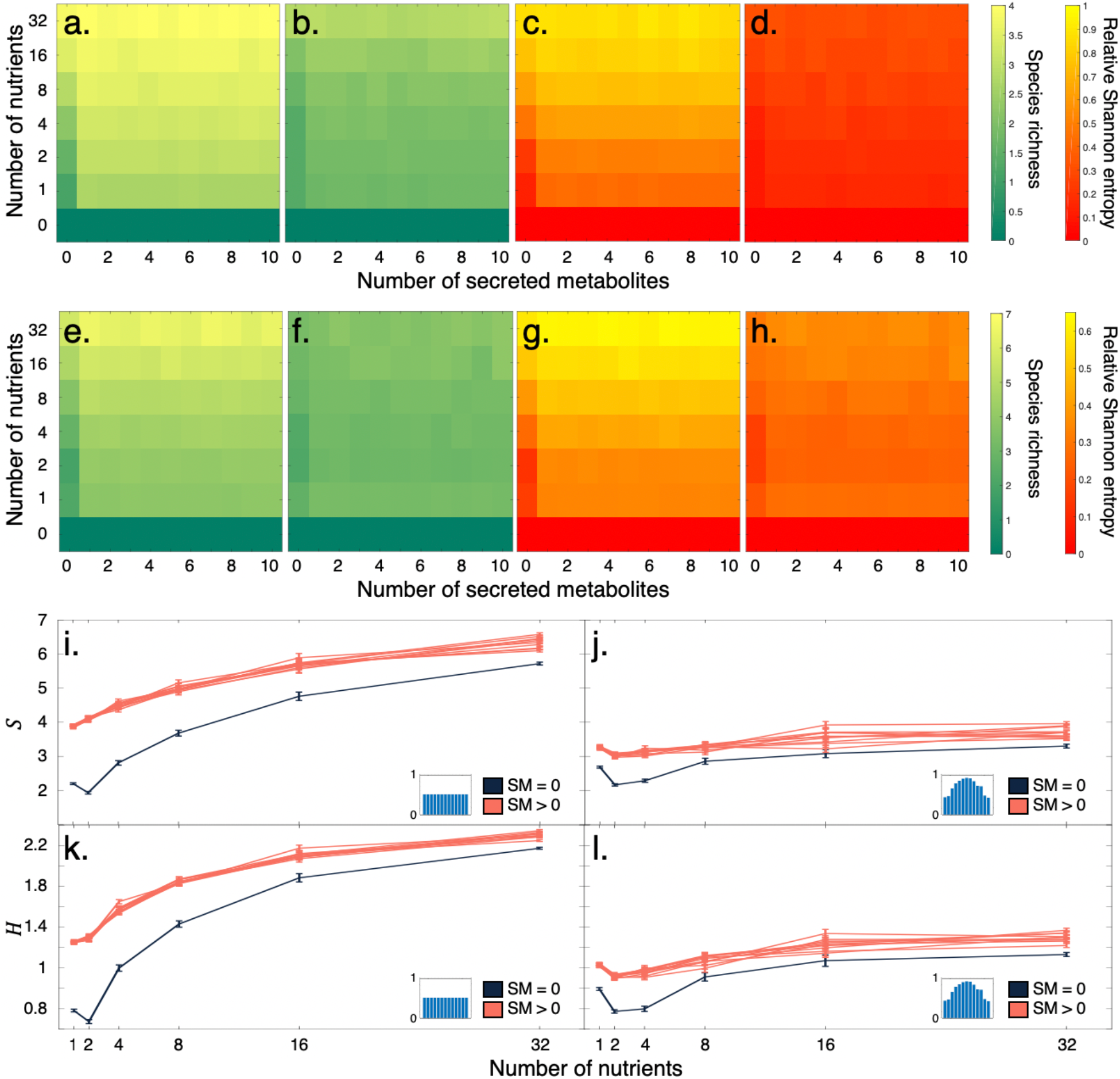
Distributions of average model-predicted species richness and Shannon entropy for simulated communities. **(a-d).** Average species richness **(a, b)** and relative Shannon entropy **(c, d)** for randomly-parametrized four-species communities containing either one **(b, d)** or no generalists **(a,c)**. **(e-h).** Average species richness **(e, f)** and relative Shannon entropy **(g, h)** for 13-species communities with either uniform **(e, g)** or experimentally-derived **(f, h)** resource utilization capabilities. Relative Shannon entropy is computed by dividing the model-predicted Shannon entropy for a given simulation by the maximum theoretical Shannon entropy for a 4- or 13-species community. **(i-l).** Distributions of species richness *S* **(i, j)** and Shannon entropy *H* **(k, l)** for 13-species communities with uniform **(i, k)** and experimentally-derived **(j, l)** nutrient utilization preferences. Each line represents the trajectory of species richness or Shannon entropy relative to the number of carbon sources supplied for a certain number (0-10) of secreted metabolites (SM) made available. Insets represent species-specific nutrient utilization probabilities. Error bars are s.e.m.

**Supplementary Figure 16.**
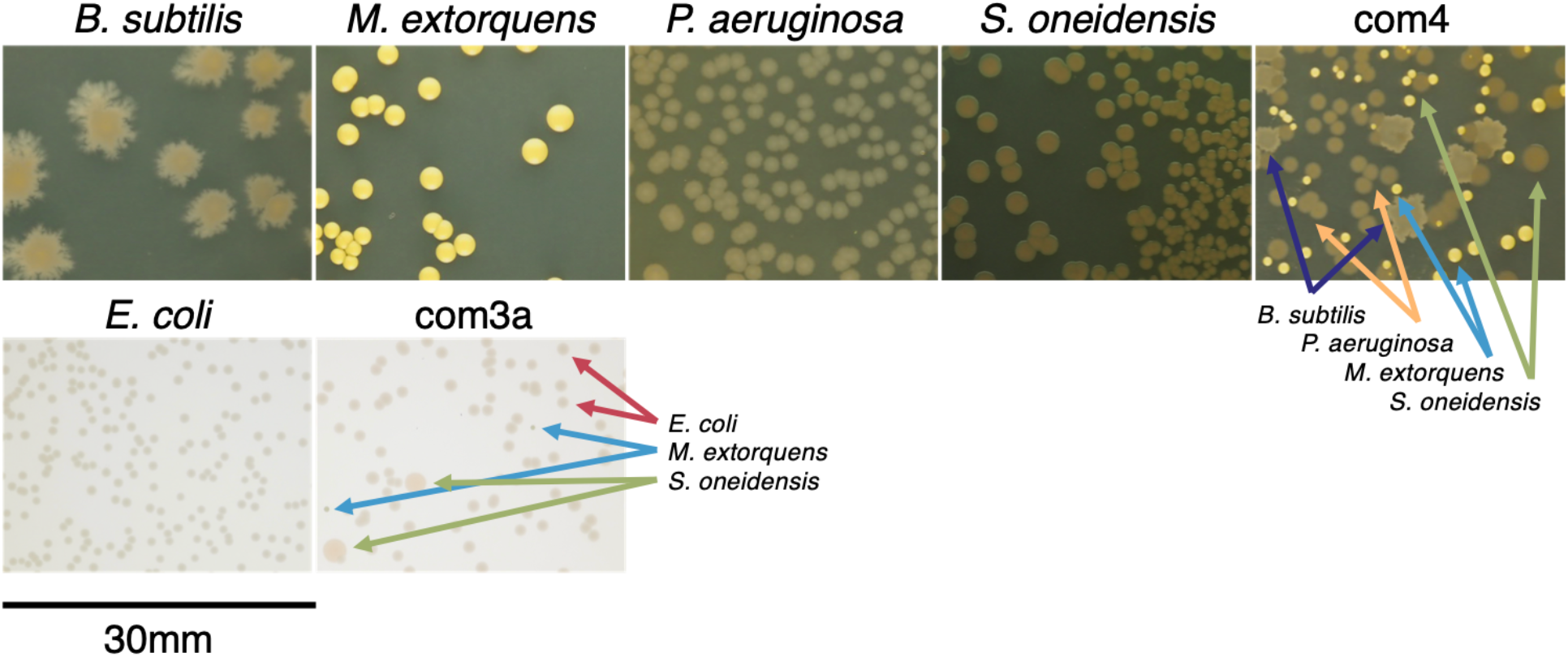
Differentiable colony morphologies for organisms assayed by agar plating.

**Supplementary Figure 17.**
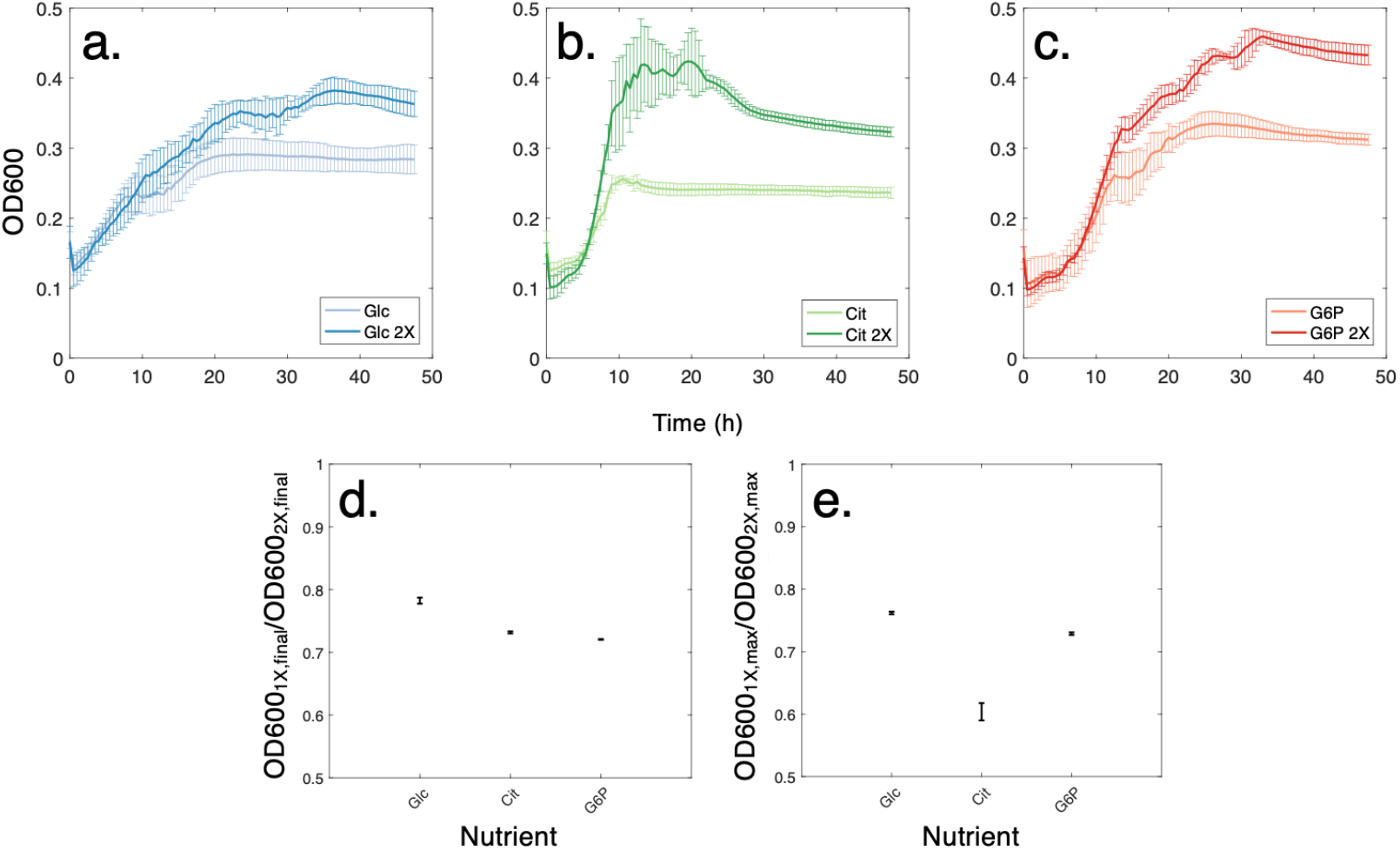
Analysis of community growth yields in single (25 mM C) and double (50 mM C) concentrations of nutrients. **(a-c).** Growth trajectories for 14-species community (com14) in two concentrations of D-glucose (Glc, **a**), citrate (Cit, **b**), and glucose-6-phosphate (G6P, **c**). **(d).** Ratio of community biomass values between single and double nutrient concentrations at endpoint **(d)** and for maximum OD values **(e)**. As the amount of resources was doubled, we expected a ratio of biomass at of 0.5 between the two concentrations. However, the observed average ratio was 0.74 ± 0.03, suggesting that some organisms grow more efficiently on diminished nutrient concentrations. Error bars indicate standard deviation.

**Supplementary Figure 18.**
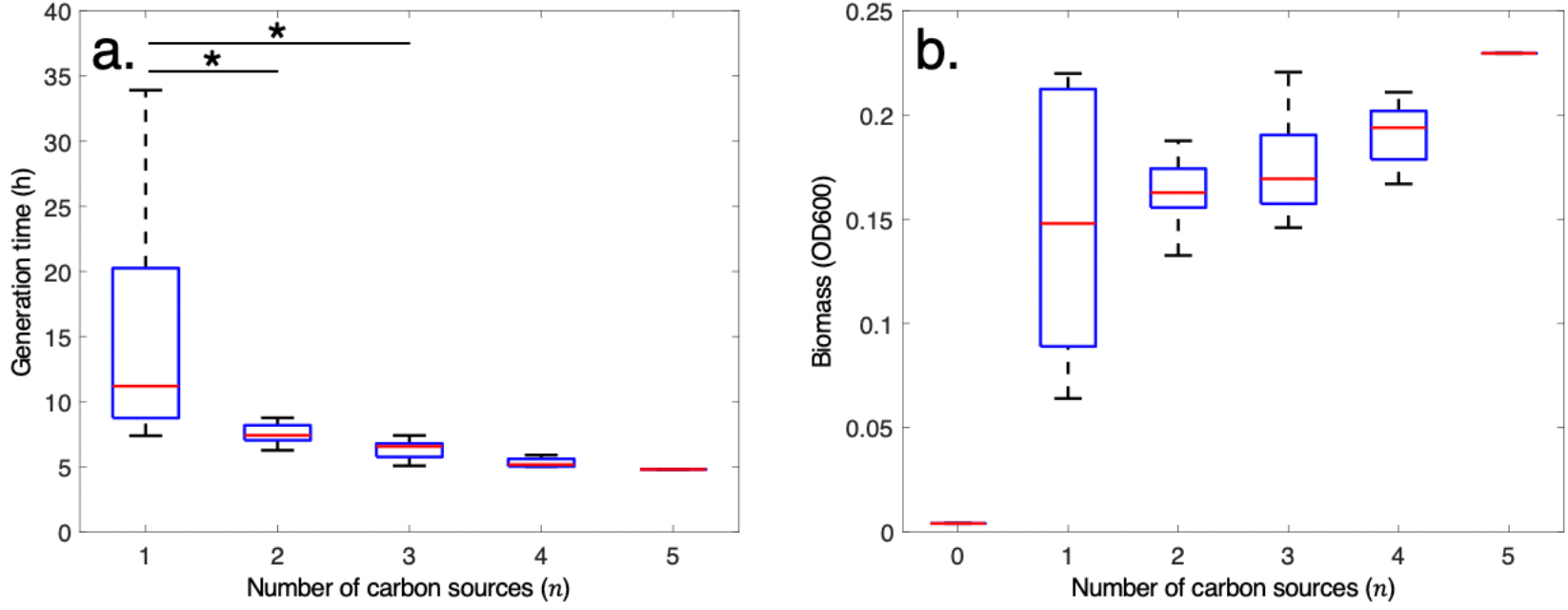
Growth phenotype for 13-species community com13a. **(a).** Generation time vs. number of nutrients for 13-species community grown on five carbon sources (com13a, Supplementary Table 5). Generation time was calculated by obtaining the maximum slope in biomass (Supplementary Figure 3b) using a moving window encompassing 5 hours. Significance is calculated by comparing generation times between single- and multiple-nutrient conditions and is indicated by (*) p < 0.05 (paired t-test *p* = 0.03 for 1 vs. 2 nutrients, 0.02 for 2 vs. 3 nutrients, 0.52 for 3 vs. 4 nutrients, and 0.33 for 4 vs. 5 nutrients). **(b).** Growth yields grouped by the number of carbon sources in each environment at the end of the experiment (144h). For complete description of sample size see Supplementary Table 4. No significant increase in yield with environmental complexity was detected (paired t-test *p* = 0.49 for 1 vs. 2 nutrients, 0.18 for 2 vs. 3 nutrients, 0.20 for 3 vs. 4 nutrients, and 0.10 for 4 vs. 5 nutrients).

**Supplementary Figure 19.**
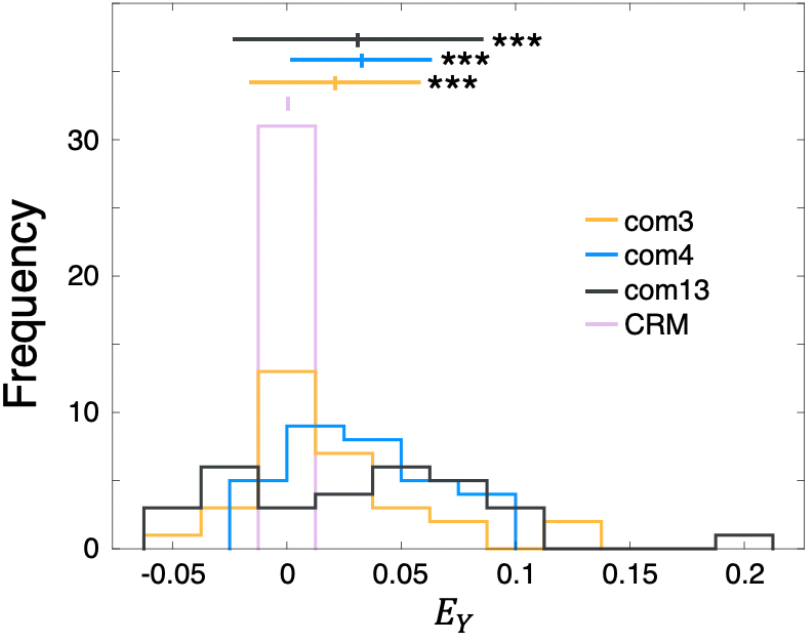
Yield epistasis *E*_*Y*_ for com3, com4, and com13 after the first 48 hours of growth. P-values for each community are: 0.002 (com3), 1.1e^−7^ (com4), and 0.001 (com13) compared against the CRM distribution.

**Supplementary Figure 20.**
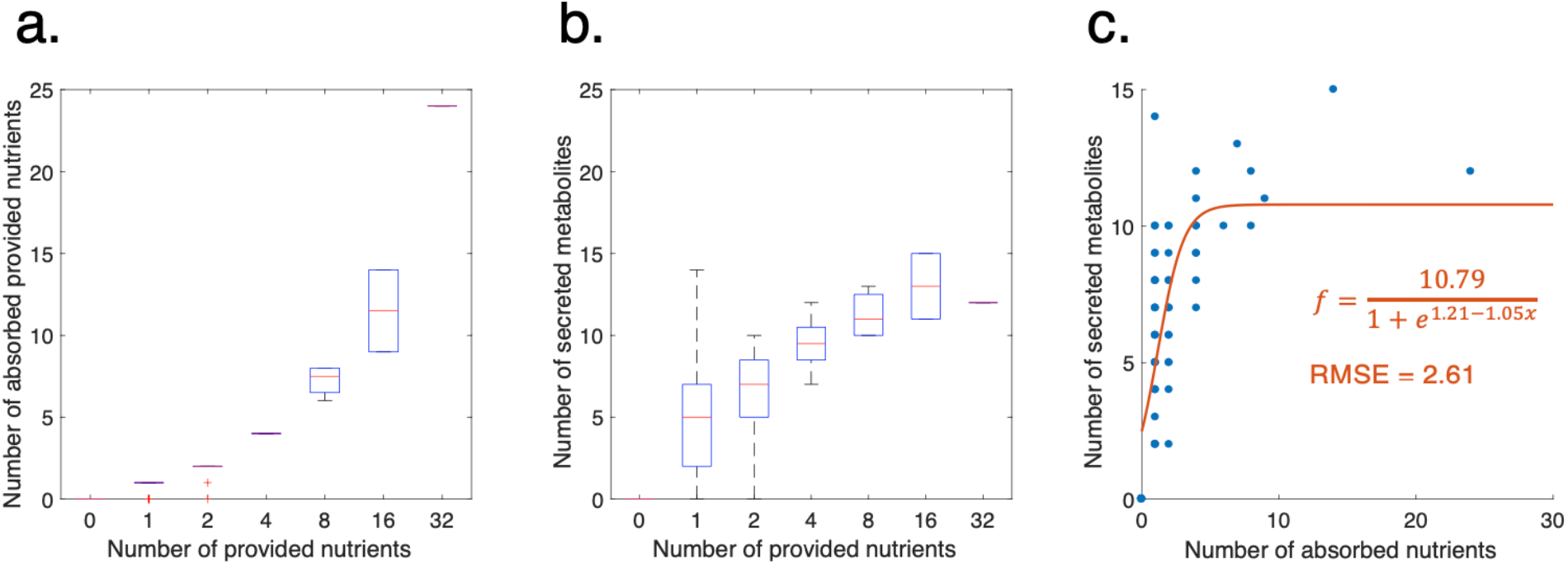
Flux balance analysis-predicted number of absorbed and secreted metabolites for 4-species community made up of *B. subtilis*, *M. extorquens*, *P. aeruginosa*, and *S. oneidensis* in combinations of up to 32 carbon sources. **(a).** Number of nutrients taken up by any of the four organisms vs. number of provided nutrients. **(b).** Number of secreted metabolites secreted by any of the four organisms vs. the number of provided nutrients. **(c).** Relationship between number of secreted metabolites and absorbed nutrients. A logistic function provided the best fit for this data, indicating a sharp rise followed by a plateau in the number of unique secreted metabolites.

**Supplementary Figure 21.**
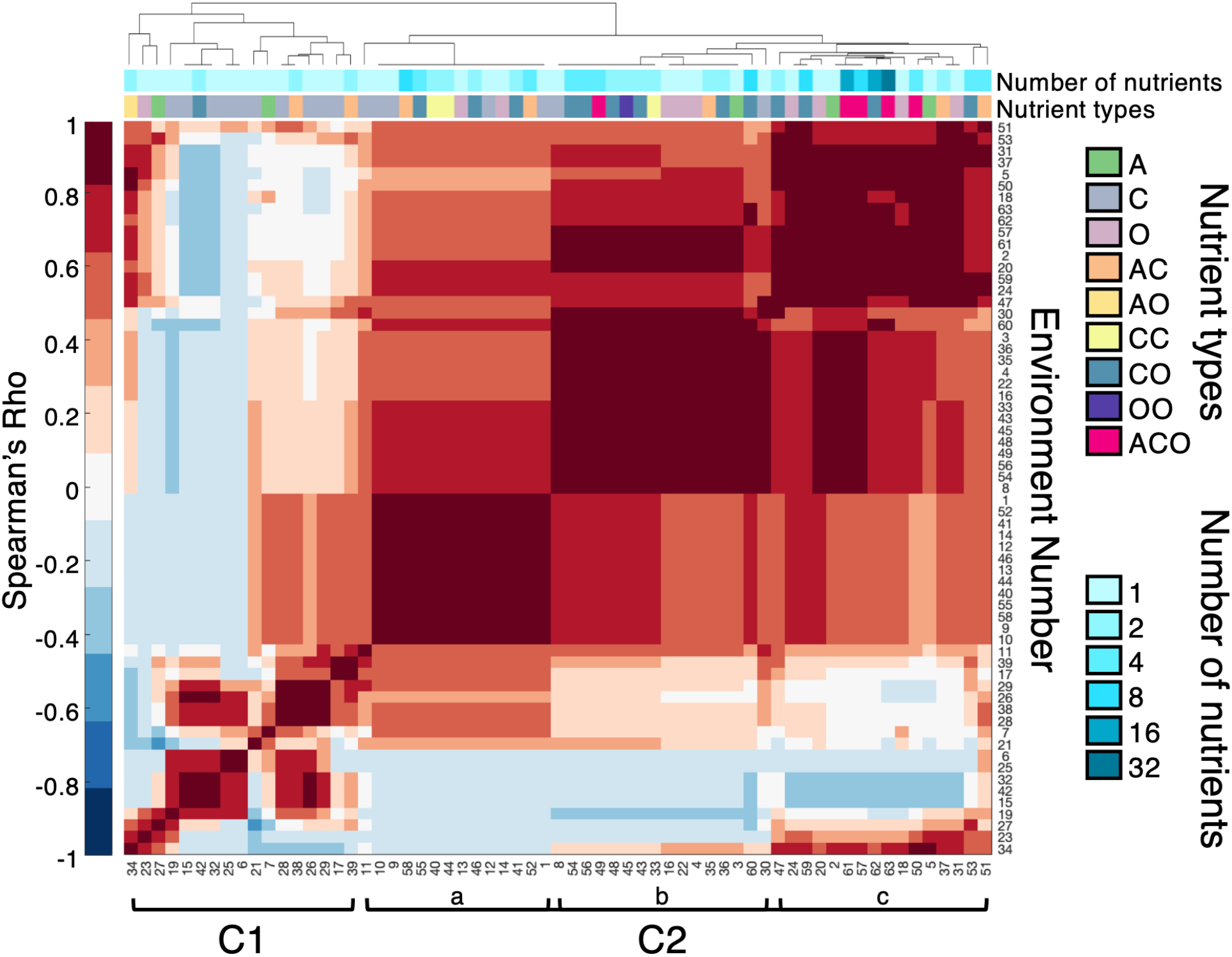
Hierarchical clustering of 63 nutrient combinations according to Spearman correlations between community taxonomic distributions. Clusters C1 and C2a-c are designated according to higher-level branches. Compositions of the 63 environments are provided in Supplementary Table 3. C1 contains 17 conditions and C2 contains the remaining 46. Conditions that were clustered closely in C1 included D-mannose and D-glcNAc + D-galacturonate (conditions 15 and 42), which displayed almost equal distributions of *E. coli* and *P. fluorescens*, as well as conditions 26, 28, 29, and 38 (D-xylose, D-galactose, D-G6P, and D-trehalose + D,L-serine), which displayed distributions of *E. coli*, *P. aeruginosa*, and *P. fluorescens*. Subcluster C2a contains a variety of nutrient combinations ranging from one to 8 carbon sources. Despite this variability, these environments all resulted in communities that were dominated by *P. aeruginosa*. Subcluster C2b was mainly represented by environments containing organic acids, which yielded communities composed of *A. baylyi* and *P. aeruginosa*. Subcluster C2c contained the most environmentally-complex conditions (with both 16-carbon source and the 32-carbon source condition) and displayed some of the most pronounced differences when compared to those in other clusters. In particular, conditions 15, 32, and 42 (D-mannose, D-sorbitol, and D-glcNAc + D-galacturonate) displayed the most dissimilar community compositions compared to subcluster C2c.

**Supplementary Figure 22.**
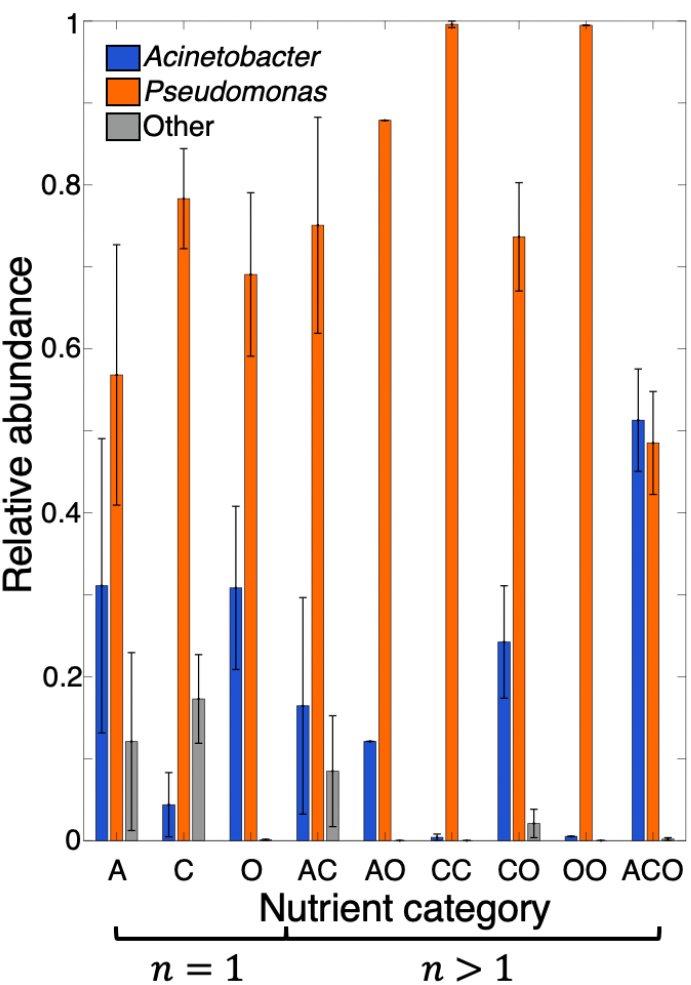
Relative abundances of organism groupings (*Acinetobacter*, *Pseudomonas*, and ‘Other’) that emerged from hierarchical clustering according to nutrient type (A: amino acid, C: carbohydrate, O: organic acid). Single nutrient conditions (*n* = 1) contain only one nutrient type, while multiple-nutrient conditions (*n* > 1) contain at least one of the nutrient types shown. For example, the L-glutamine condition would be categorized under ‘A’ for amino acid, while the L-glutamine + formate condition would be categorized as ‘AO’ as it contains both an amino acid and an organic acid. Nutrient-specific type designations are provided in Supplementary Table 2. Error bars indicate standard deviation.

**Supplementary Figure 23.**
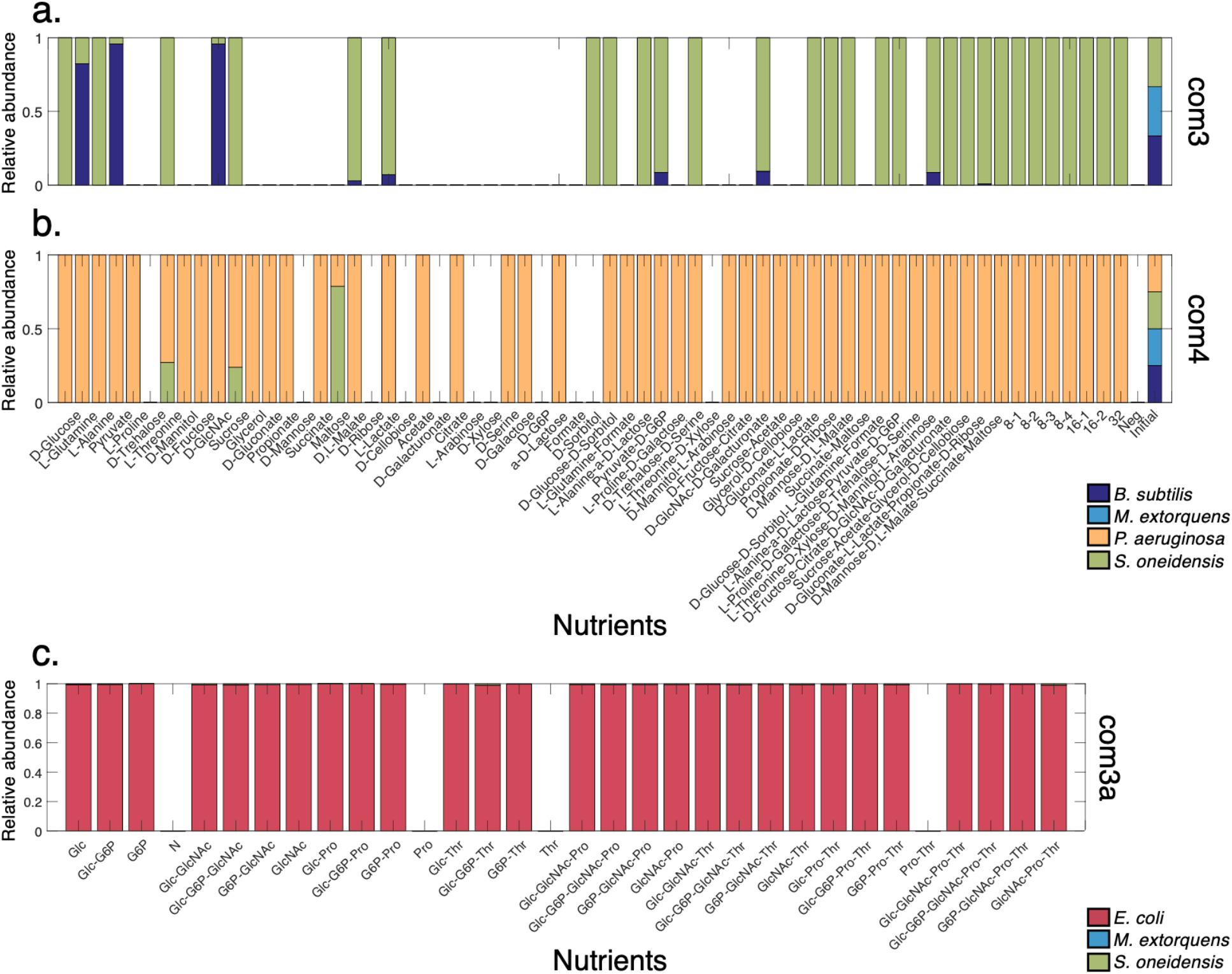
Endpoint species relative abundances for multispecies communities com3 **(a)** and com4 **(b)**, and com3a **(c)**. *P. aeruginosa* was most often dominant in com4, outcompeting the other community members in 52 cases. In addition to the dominance of *P. aeruginosa* in com4, we noticed that *S. oneidensis* was dramatically overrepresented in our com3 experiment. This distribution was striking, as we expected *M. extorquens* to be most often dominant given its wider breadth of nutrient utilization capabilities in monoculture (Supplementary Figure 2a). **(c)**. An additional experiment containing *E. coli*, *M. extorquens*, and *S. oneidensis* (com3a, Supplementary Table 5) similarly highlighted the ability of a single organisms to overtake small communities. Relative abundances for com3 and com4 are adjusted based on a calibration of CFU counts to equal OD600 values. Unabbreviated environmental compositions for com3 and com4 are outlined in Supplementary Table 3. Unique conditions for com3a are: D-glucose (DGlc), pyruvate (Pyr), GlcNAc (GlcNAc), L-proline (Pro), L-threonine (Thr), and a no-carbon negative control (N). Missing bars indicate no growth in the specified condition.

**Supplementary Figure 24.**
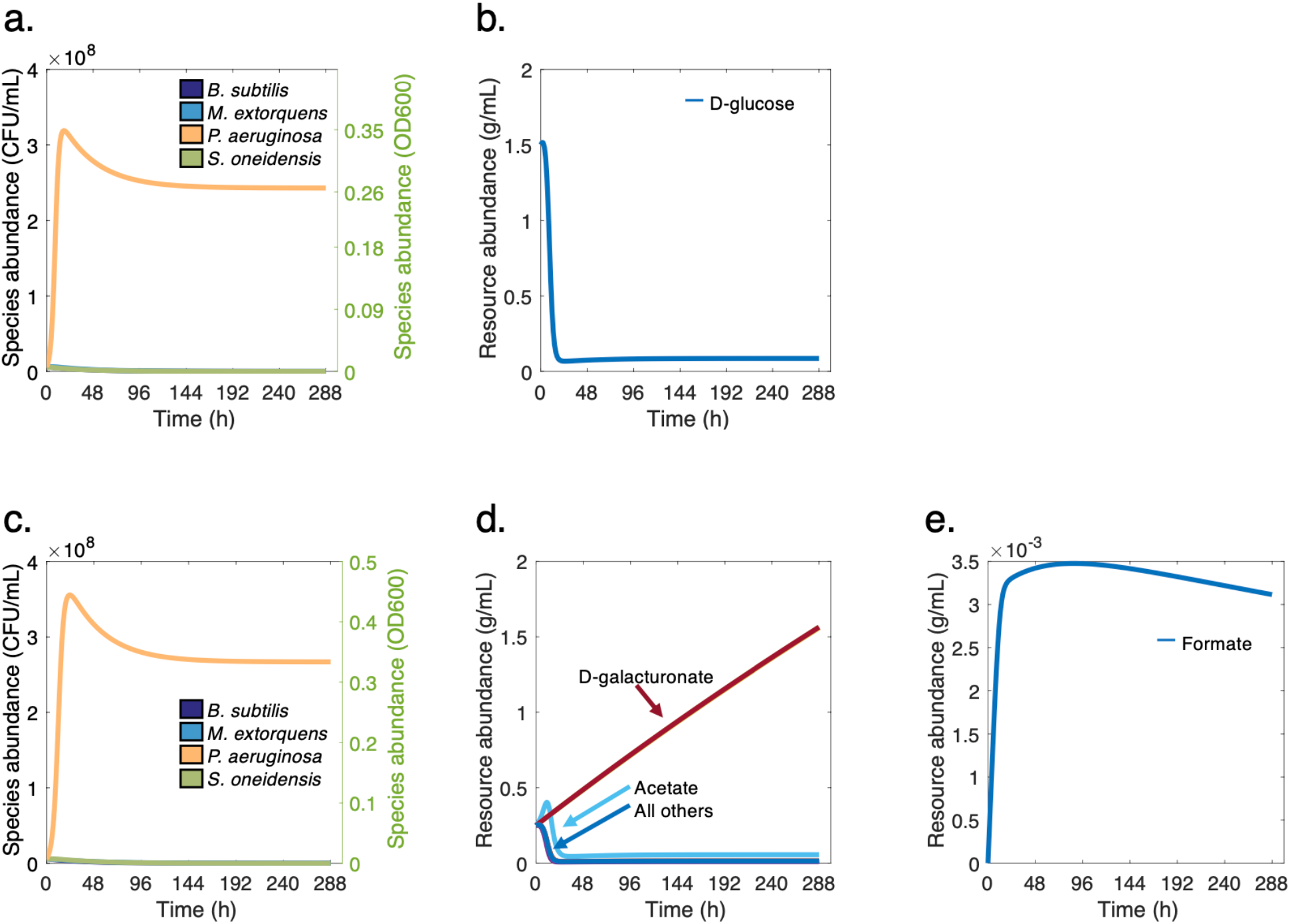
Example consumer resource model-predicted growth phenotypes for com4 grown on D-glucose **(a)** and D-fructose, citrate, D-glcNAc, D-galacturonate, sucrose, acetate, glycerol, and D-cellobiose **(b)**, (conditions 1 and 59, respectively, Supplementary Table 3). Community compositions and yields recapitulate those observed experimentally (Supplementary Figure 7a, Supplementary Figure 21a) and reveal environment-specific nutrient utilization patterns. Specifically, the community grown on the second condition shown is predicted to consume all of the provided nutrients except D-galacturonate, which accumulates in the medium. Moreover, the community is predicted to first rapidly secrete then stop producing formate, which becomes diluted over time.

**Supplementary Figure 25.**
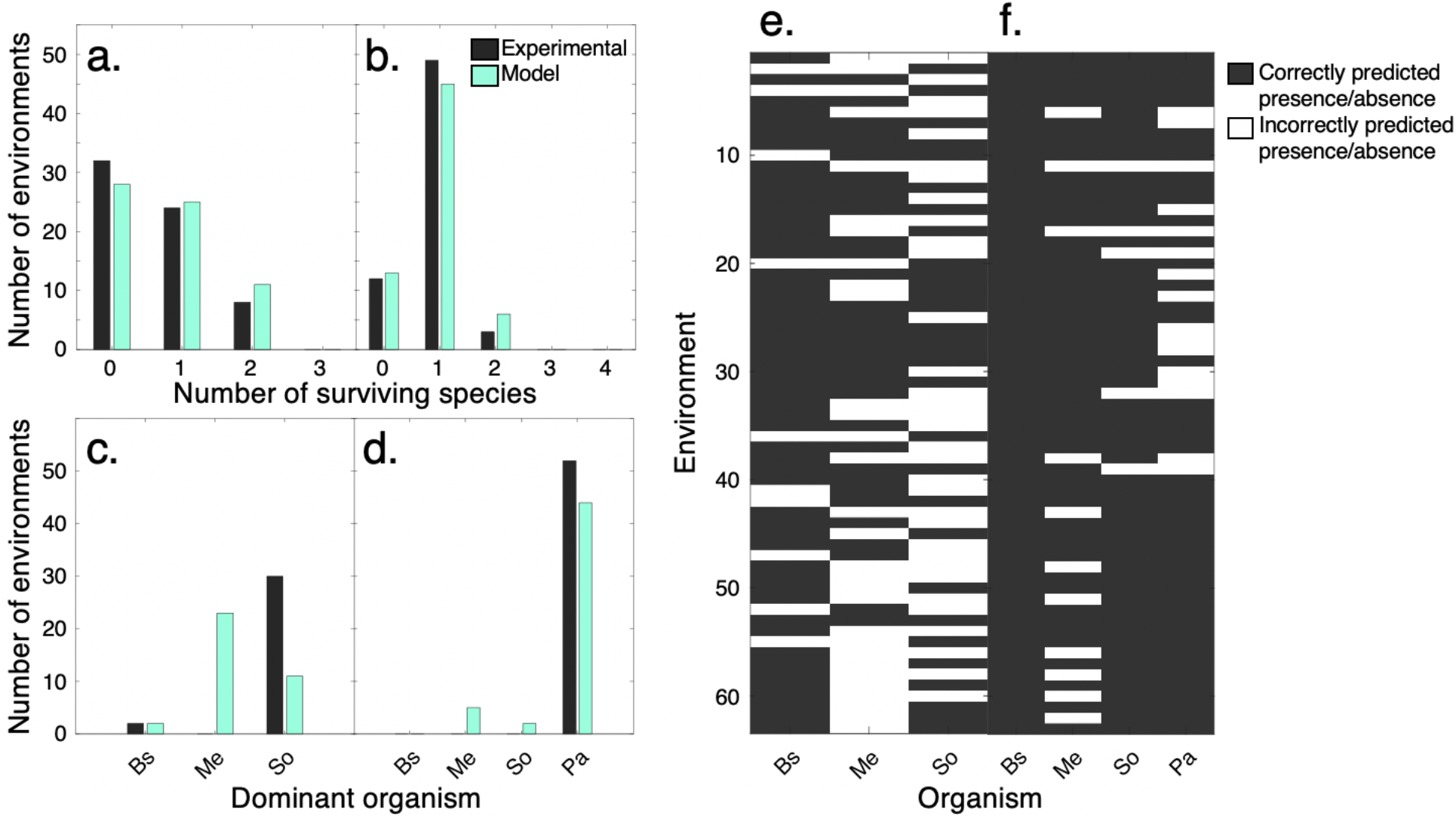
Consumer resource model predictions compared to com3 and com4 experiments. **(a, b).** Frequency of number of environments by number of surviving species for com3 **(a)** and com4 **(b)** as observed experimentally (gray) and predicted by the consumer resource model (light blue). **(c, d).** Frequency of environments in which each organism was predicted to be dominant in com3 **(c)** and com4 **(d)**, as observed experimentally (gray) and predicted by the consumer resource model (light blue). Organisms are abbreviated as follows: Bs: *B. subtilis*, Me: *M. extorquens*, So: *S. oneidensis*, and Pa: *P. aeruginosa*. **(e, f).** Environment-by-environment comparison of model-predicted and experimentally-observed presence/absence of individual organisms in com3 **(e)** and com4 **(f)**. Species-specific accuracies are: 84.1% for Bs, 52.4% for Me, and 47.6% for So in com3, and 100% for Bs, 82.5% for Me, 92.1% for So, and 74.6% for Pa in com4.

### Supplementary Tables

**Supplementary Table 1.** Complete list of organisms used in experiments.

| Organism name | Strain | ATCC | Temp.<br>(°C) | Flux Balance<br>Model |
| --- | --- | --- | --- | --- |
| *Acinetobacter baylyi* | ADP1 | 33305 | 37 | 55 |
| *Bacillus licheniformis* | 46 | 14580 | 37 |  |
| *Bacillus subtilis* | 168 | 23857 | 37 | 42 |
| *Corynebacterium glutamicum* | 534 | 13032 | 37 |  |
| *Escherichia coli* | MG1655 | 25922 | 37 | 56 |
| *Lactococcus lactis* | MG1363 | 19257 | 37 | 57 |
| *Methylobacterium extorquens* | AM1 | 43645 | 30 | 43 |
| *Pseudomonas aeruginosa* | PA01 | 9027 | 37 | 44 |
| *Pseudomonas fluorescens* | 28/5 | 13525 | 26 |  |
| *Pseudomonas putida* | KT2440 | 47054 | 26 | 58 |
| *Streptomyces coelicolor* | A3(2) | 23899 | 28 | 59 |
| *Shewanella oneidensis* | MR-1 | 700550 | 30 | 45 |
| *Salmonella enterica* | LT2 | 27106 | 37 | 60 |
| *Streptococcus thermophilus* | LMG18311 | BAA-250 | 37 |  |
| *Vibrio natriegens* | 111 | 14048 | 26 |  |

**Supplementary Table 2.**
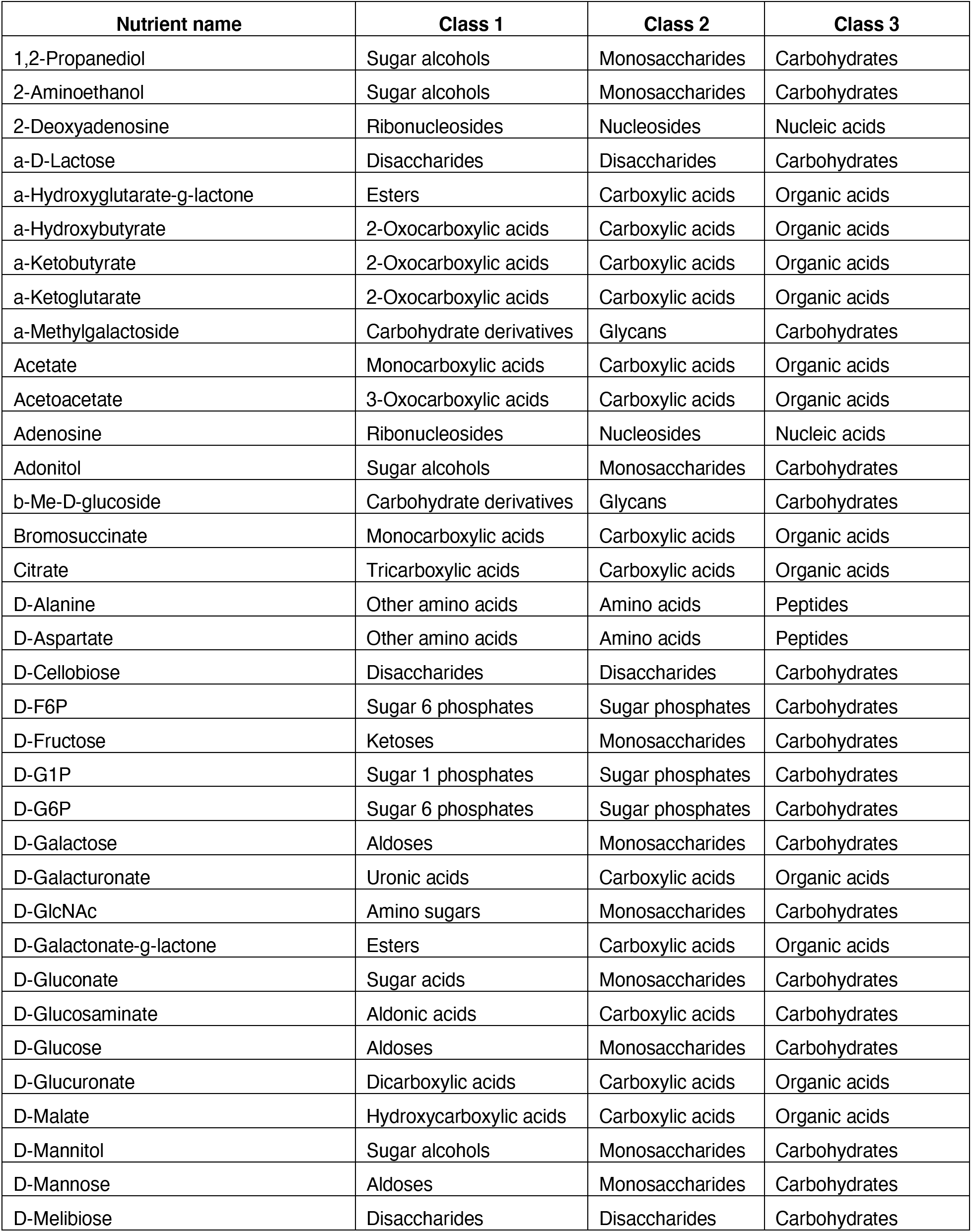
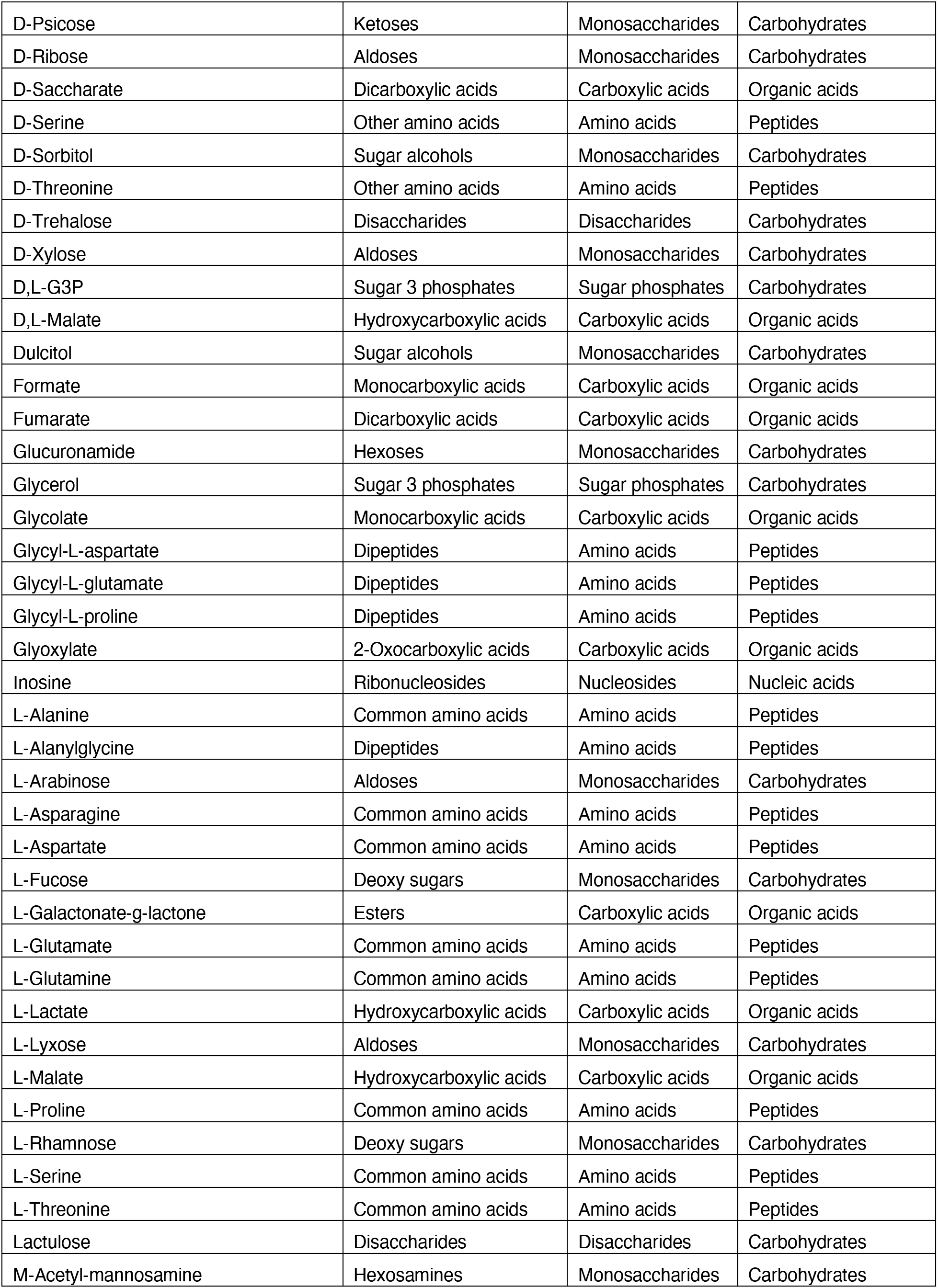
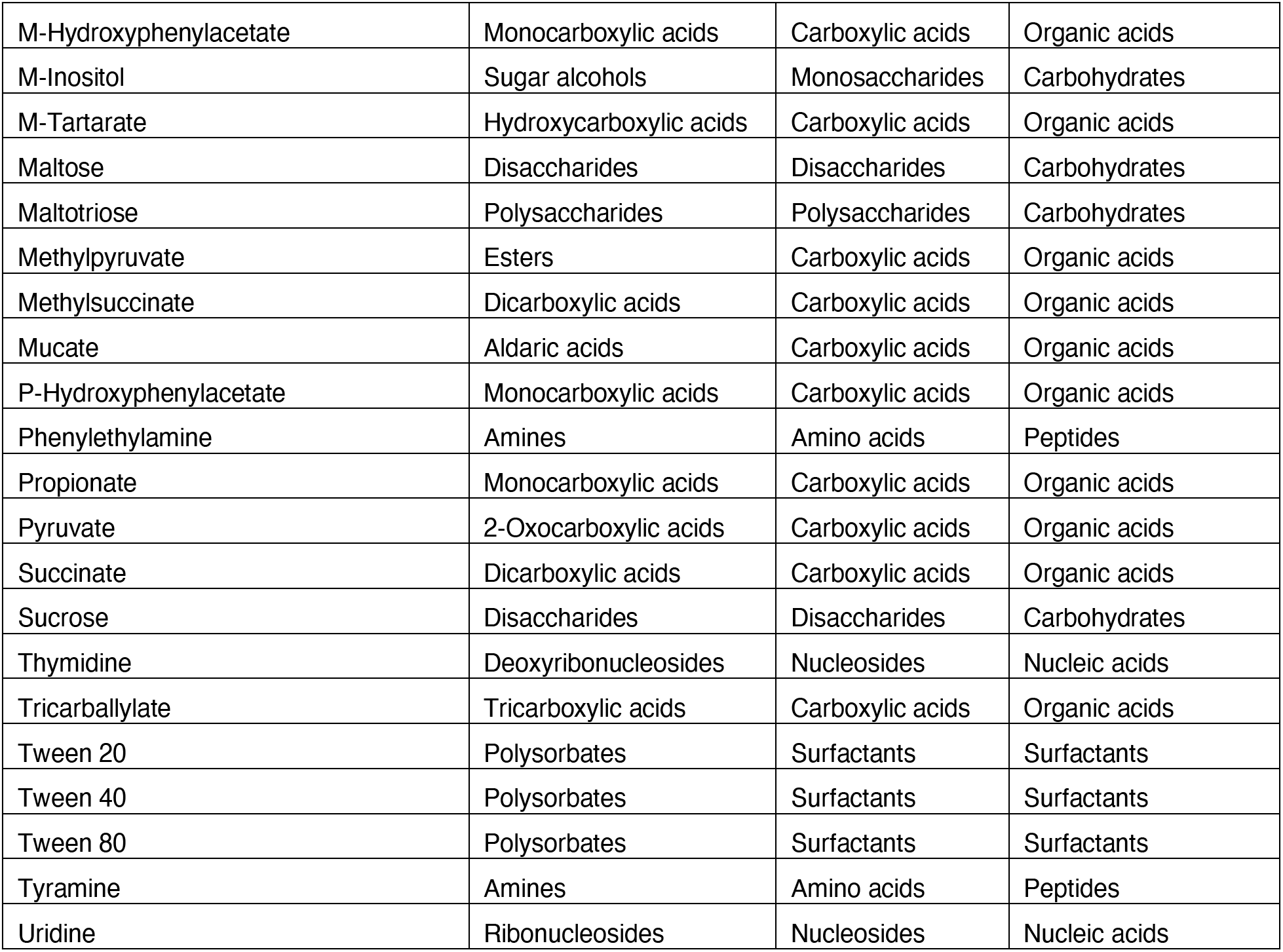
List and ontology of 95 carbon sources in Biolog PM1 plate.

**Supplementary Table 3.**
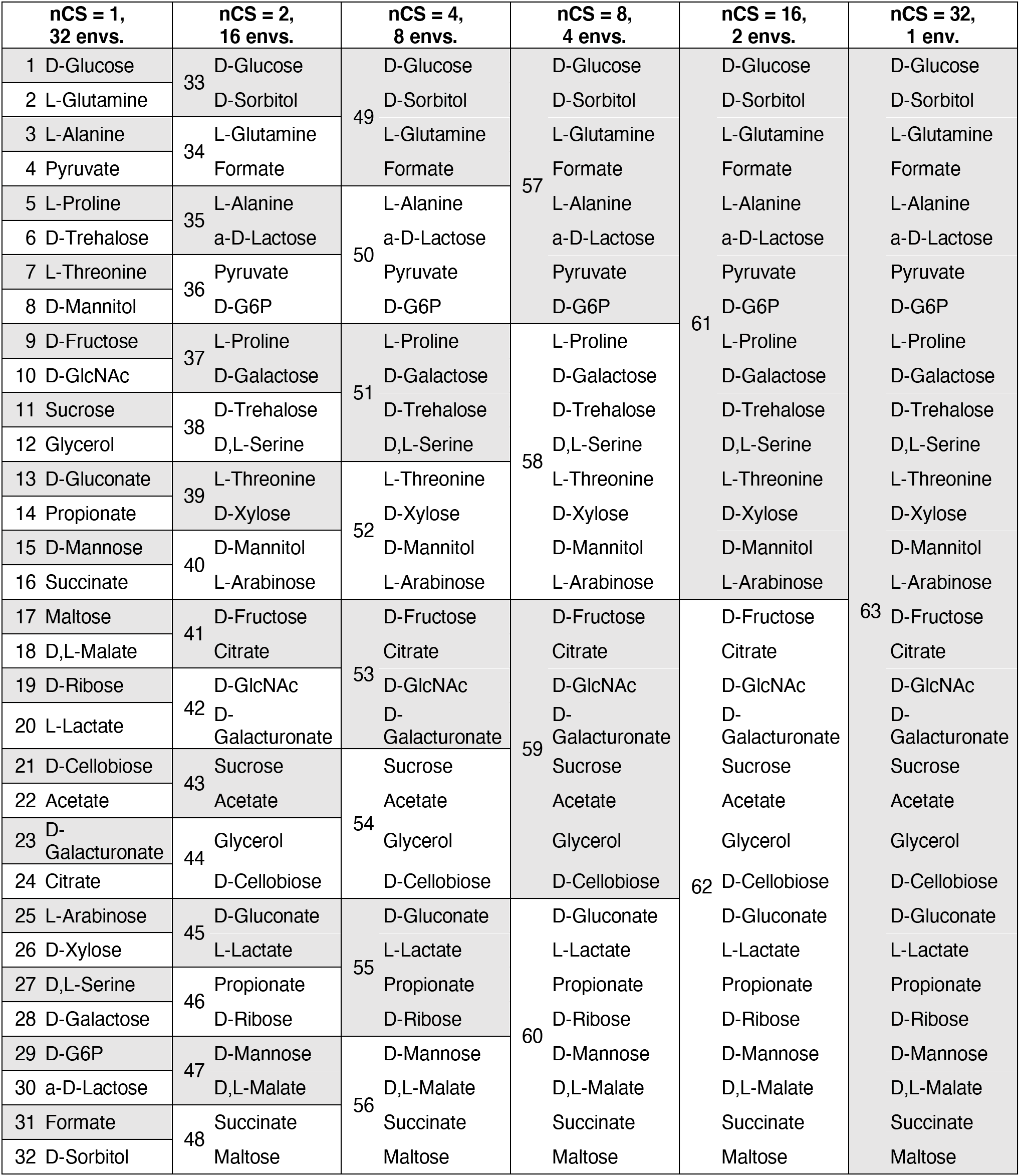
Nutrient pairings in 32-carbon source experiments (com3, com4, com13). Each individual nutrient combination is numbered and highlighted with alternating background colors, for a total of 63 unique conditions containing varying numbers of carbon sources (nCS).

**Supplementary Table 4.**
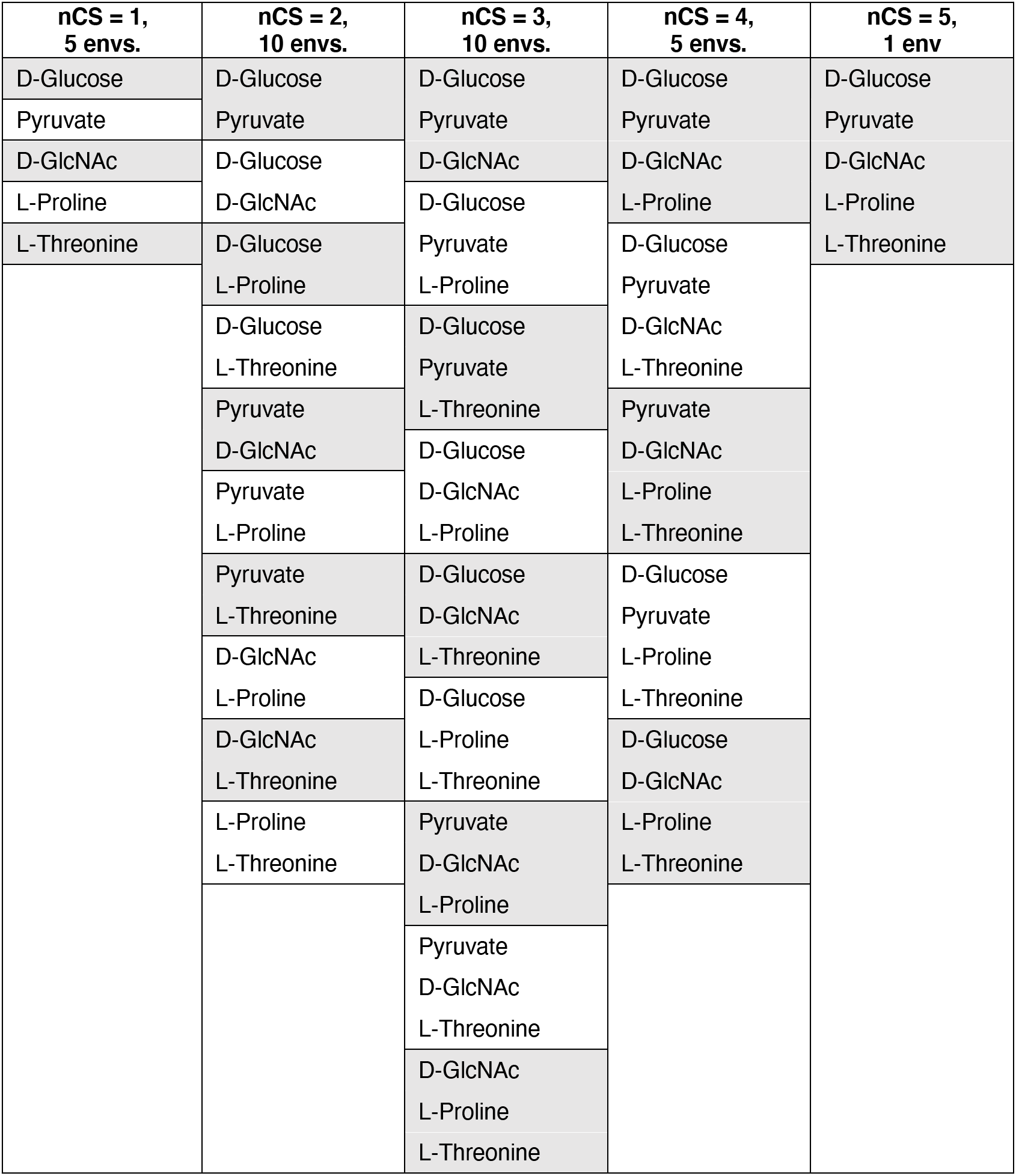
Nutrient pairings in 5-carbon source experiments (com3a, com13a). Each background color indicates one environmental composition, for a total of 31 unique conditions containing varying numbers of carbon sources (nCS).

**Supplementary Table 5.** Descriptions of combinatorial nutrient experiments. All experiments were carried out at 30°C and provided 50 mM C per well (com14 also grown at 25 mM C).

| Name | Number of organisms | Organisms | Number of nutrients | Number of passages | Passaging frequency |
| --- | --- | --- | --- | --- | --- |
| com3 | 3 | *B. subtilis*, *M. extorquens*, *S. oneidensis* | 32 (Supplementary Table 3) | 6 | 48h |
| com3a | 3 | *E. coli*, *M. extorquens*, *S. oneidensis* | 5 (Supplementary Table 4) | 0 (24h batch) | 0 |
| com4 | 4 | com3 + *P. aeruginosa* | 32 (Supplementary Table 3) | 6 | 48h |
| com13 | 13 | com4 + *A. baylyi*, *B. licheniformis*, *C. glutamicum*, *E. coli*, *L. lactis*, *P. fluorescens*, *P. putida*, *S. coelicolor*, *V. natriegens* | 32 (Supplementary Table 3) | 6 | 48h |
| com13a | 13 | com13 | 5 (Supplementary Table 4) | 6 | 24h |
| com14 | 14 | com13 + *S. enterica* | 4 (Supplementary Figure 3a) | 0 (24h batch) | 0 |

**Supplementary Table 6.** Regression coefficients for individual carbon sources significantly positively or negatively associated with higher Shannon entropy in com13. We generated a simple linear regression model that relates the presence of each carbon source to the Shannon entropy exhibited by the communities. We calculated regression coefficients for each nutrient, allowing us to estimate the contribution of each carbon source to taxonomic balance independent of the number of nutrients. Here, we estimated D-galactose to be the most highly associated with greater community evenness. Indeed, the environment containing only D-galactose yielded a community with a relatively even composition of *E. coli*, *P. aeruginosa*, and *P. fluorescens*, while the community grown in the two-carbon-source condition with D-galactose also contained three organisms (*A. baylyi*, *P. aeruginosa*, and *P. putida)* (Figure 3a). Despite these relatively balanced communities at lower complexities, the presence of a nutrient significantly associated with lower Shannon entropy like L-arabinose can overpower the effects of nutrients like D-galactose. This effect is most clearly observed in the 8-carbon source condition containing both these nutrients, which resulted in the complete dominance of *P. aeruginosa*.

| Nutrient | Coefficient | P-value |
| --- | --- | --- |
| L-Glutamine | 0.619 | 2.78E-02 |
| D,L-Malate | 0.858 | 2.40E-03 |
| D-Ribose | 0.795 | 4.90E-03 |
| D-Cellobiose | 1.049 | 2.00E-04 |
| Citrate | 1.049 | 2.00E-04 |
| D-Xylose | 0.673 | 1.69E-02 |
| D-Galactose | 0.752 | 7.70E-03 |
| D-Glucose | -0.826 | 3.50E-03 |
| D-Fructose | -0.951 | 8.00E-04 |
| Glycerol | -1.285 | 0.00E+00 |
| D-Gluconate | -0.872 | 2.10E-03 |
| Propionate | -1.205 | 0.00E+00 |
| L-Arabinose | -1.105 | 1.00E-04 |

**Supplementary Table 7.** Minimal medium composition for flux-balance modeling.

| Metabolite | BIGG metabolite name<sup>61</sup> |
| --- | --- |
| Calcium | ca2[e] |
| Chloride | cl[e] |
| Cobalt | cobalt2[e] |
| Copper<sup>2+</sup> | cu2[e] |
| Iron<sup>2+</sup> | fe2[e] |
| Iron<sup>3+</sup> | fe3[e] |
| Hydrogen | h2[e] |
| Water | h2o[e] |
| Potassium | k[e] |
| Magnesium | mg2[e] |
| Manganese | mn2[e] |
| Molybdate | mobd[e] |
| Sodium | na1[e] |
| Ammonium | nh4[e] |
| Nickel<sup>2+</sup> | ni2[e] |
| Nitrate | no3[e] |
| Oxygen | o2[e] |
| Phosphate | pi[e] |
| Sulfate | so4[e] |
| Zinc<sup>2+</sup> | zn2[e] |

**Supplementary Table 8.** List of unique organic molecules predicted to be secreted across all flux-balance simulations.

| Metabolite | BIGG metabolite name<sup>61</sup> |
| --- | --- |
| Acetate | ac[e] |
| D-alanine | ala-D[e] |
| Allantoin | alltn[e] |
| Citrate | cit[e] |
| Formate | for[e] |
| Formamide | frmd[e] |
| Fumarate | fum[e] |
| Glycolaldehyde | gcald[e] |
| D-gluconate | glcn-D[e] |
| Glycolate | glyclt[e] |
| L-lactate | lac-L[e] |
| D-malate | mal-D[e] |
| R-pantothenate | pnto-R[e] |
| Riboflavin | ribflv[e] |
| Spermidine | spmd[e] |
| Succinate | succ[e] |
| Trehalose | tre[e] |
| Urea | urea[e] |

**Supplementary Table 9.** Descriptions and quantities for consumer resource model state variables and parameters. Resource supply and dilution rates are defined by 48-hour experimental timescale.

| Variable/parameter | Description | Units | Quantity |
| --- | --- | --- | --- |
| $N_i$ | Abundance of organism $i$ | CFU/mL | variable |
| $R_\alpha$ | Abundance of nutrient $\alpha$ | g/mL | variable |
| $C_{i\alpha}$ | Uptake rate per unit concentration of resource $\alpha$ by organism $i$ | mL/hr | variable |
| $D_{\alpha\beta i}$ | Proportion of nutrient $\alpha$ converted to nutrient $\beta$ by organism $i$ | unitless | variable |
| $l_\alpha$ | Leakage fraction for resource $\alpha$ | unitless | 0.25 [18] |
| $g_i$ | Conversion factor from energy uptake to growth rate for organism $i$ | 1/energy | 1 [18] |
| $w_\alpha$ | Energy content of resource $\alpha$ | energy/g | $1 \times 10^8$ |
| $k_{i,a}$ | Half velocity constant for resource uptake | g/mL | 3000 |
| $m_i$ | Minimal energy uptake for maintenance of each species $i$ | energy/hr | 0.05 |
| $\kappa_\alpha$ | External supply of resource $\alpha$ | g/mL/hr | $\frac{1.5g/mL}{48h}$ |
| $d$ | Dilution rate | 1/hr | $\left(\frac{10\mu l}{300\mu l}\right) \times 48h^{-1}$ |

## Notes

### Competing Interest Statement

The authors have declared no competing interest.

